# Assembling Ruthenium Complexes to Form Ruthenosome Unleashing Ferritinophagy-mediated Tumor Suppression

**DOI:** 10.1101/2024.10.22.619580

**Authors:** Caiting Meng, Shuaijun Li, Yana Ma, Hongwen Yu, Jiaqi Song, Junchao Zhi, Bin Zhu, Liang Shao, Xinling Liu, Lulu Yang, Mingzhen Zhang, Ye Zhang, Guanying Li

## Abstract

We introduce ruthenosomes, a fusion of liposomal and reactive oxygen species (ROS)–generating properties meticulously engineered as potent ferroptosis inducers (FINs), marking a significant advancement in metallodrug design for cancer therapy. Formed through the self-assembly of oleate-conjugated ruthenium complexes, these ruthenosomes exhibit exceptional cellular uptake, selectively accumulating in mitochondria and causing substantial disruption. This targeted mitochondrial damage significantly elevates ROS levels, triggering autophagy and selectively activating ferritinophagy. Together, these processes sensitize cancer cells to ferroptosis. In vivo, ruthenosomes effectively suppress colorectal tumor growth, underscoring their therapeutic potential. Our study pioneers a design strategy that transforms ruthenium complexes into liposome-like structures capable of inducing ferroptosis independent of light activation. By leveraging ruthenosomes as multifunctional nanocarriers, this research offers a versatile and powerful platform for ROS-mediated, ferroptosis-driven cancer cell eradication.

## Introduction

While traditional chemotherapeutic drugs are successful at inducing apoptosis in cancer cells, their efficacy is often limited by the inherent constraints of apoptotic pathways and the frequent emergence of drug resistance.^1^ This has driven growing interest in alternative cell death mechanisms as potential strategies for cancer treatment. Among these, ferroptosis—a non-apoptotic form of regulated cell death (RCD)–has shown significant promise. Ferroptosis is intricately driven by iron-dependent lipid peroxidation, making cancer cells, which often have distinct metabolic profiles and elevated reactive oxygen species (ROS) levels, particularly vulnerable to this form of cell death.^2^ Additionally, key tumor suppressors have been identified as modulators of ferroptosis, highlighting its potential to inhibit cancer progression. These unique molecular mechanisms position ferroptosis inducers (FINs) as powerful tools for overcoming chemotherapy resistance, offering a promising therapeutic strategy for treating refractory cancers.^3^

Ferroptosis can be pharmacologically targeted at multiple stages, with FINs broadly classified into three main categories: iron-based molecules or nanocomposites that target iron metabolism and induce intracellular iron overload; glutathione (GSH)-depleting agents that disrupt the ferroptosis defence system by increasing ROS levels,^4,5^ and inhibitors or modulators of ferroptosis-related enzymes^4, 5^ that covalently bind to redox homoeostasis regulators like glutathione peroxidase 4 (GPX4) to inhibit its enzymatic activity;^6^ ultimately leading to cell death. However, the efficacy of these FINs varies significantly across different cancer types, primarily due to tumor heterogeneity, which complicates the therapeutic targeting of ferroptosis and poses a persistent challenge in developing effective treatments.^7^ While excessive ROS are generally regarded as universal toxins, the direct induction of the lipid peroxidation through ROS production may offer a promising strategy for inducing ferroptosis across a wide range of cancer types.^8, 9^ Nevertheless, the non-specific nature of ROS production presents a significant challenge, as excessive ROS can activate other forms of cell death, including autophagy,^10^ apoptosis,^11, 12^ and necroptosis.^13^ This lack of specificity complicates the development of ROS generators as FINs.

A critical challenge in designing effective ROS generators for ferroptosis is the efficient production of highly reactive hydroxyl radicals (HO^•^) through the Fenton reaction, which are essential for driving the ferroptotic process. The precursors (hydrogen peroxide, H_2_O_2_)^14^ and catalysts (free Fe^2+^) required for Fenton chemistry are co-localized within the mitochondrial matrix,^15^ making mitochondria the primary site for sustained HO• generation and a key target for ferroptosis regulation.^16^ However, exploiting mitochondrial vulnerability to effectively induce ferroptosis remains challenging, with only a limited number of studies addressing this aspect in the context of tumor suppression.^17^ Given the demand in balancing the specificity and efficacy of ROS production–essential for overcoming the hurdles posed by tumor heterogeneity and harnessing ferroptosis as a therapeutic strategy, we reported here the design of a ruthenium complex assembled liposome, termed ruthenosome, that selectively bursts in ROS production in mitochondria to induce lipid peroxidation for ferroptosis-mediated cell death, demonstrating broad applicability across various cancer types (Figure 1).

**Figure 1.**
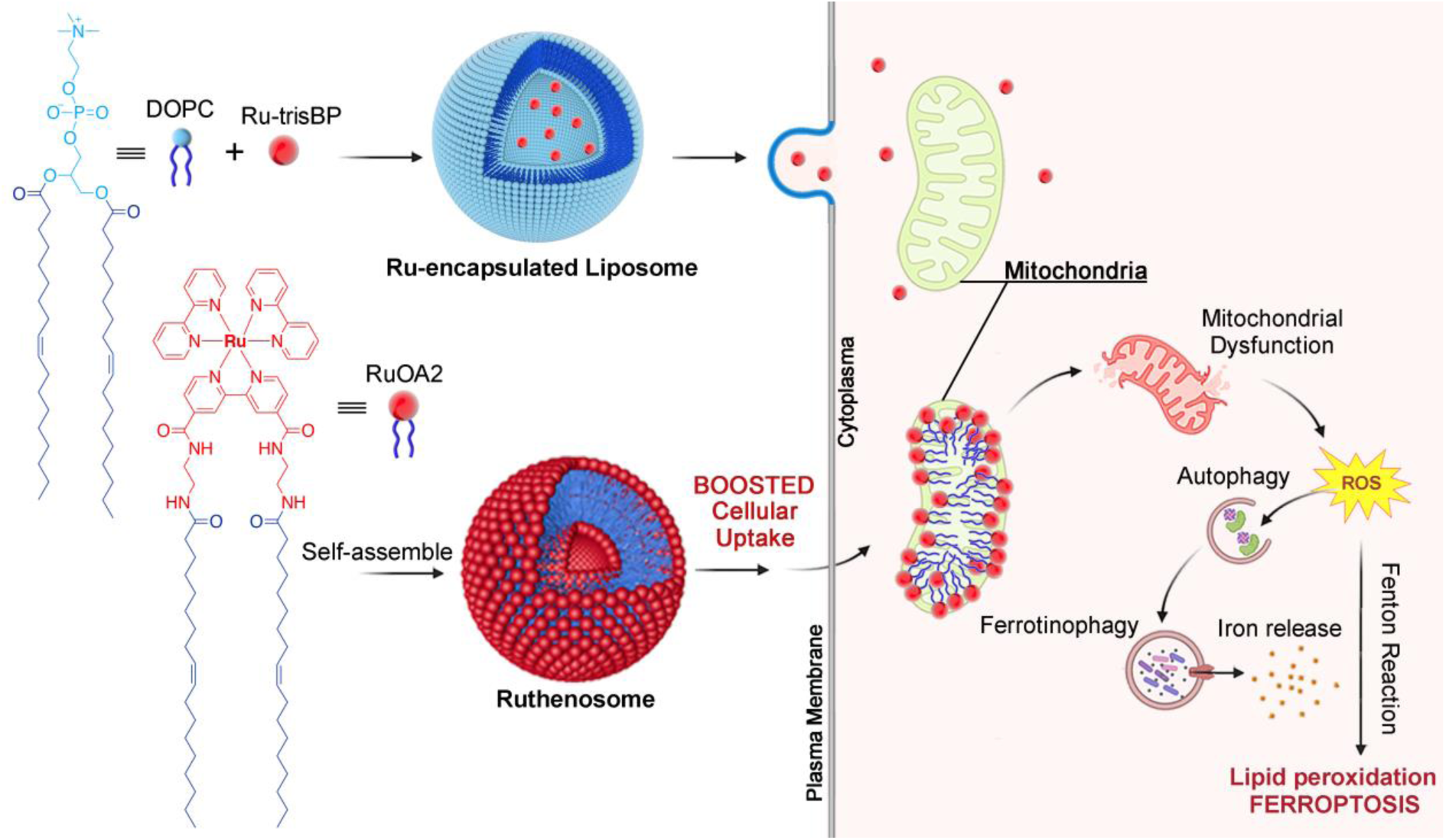
Schematic illustration of ruthenosome. Formulations of Ru-trisBP-encapsulated liposomes and **RuOA2**-assembled ruthenosomes. The ruthenosome boosts the cellular uptake of **RuOA2**, specifically targeting mitochondira and generating ROS to induce ferroptosis.

## Results

### Construct active ruthenosome via dynamic self-assembly of RuOA2

Ruthenium(II) complexes have been extensively utilized as ROS generators through mitochondrial targeting^18^ or light irradiation^19^, inducing apoptosis in cancer cells. However, their potential as FINs is often limited by inadequate cellular uptake/accumulation and insufficient ROS generation. A recent study achieved a significant breakthrough by developing a Ru(II) polypyridine complex that exhibited enhanced cellular uptake and increased ROS production, ultimately leading to photoinduced ferroptosis in breast cancer cells,^20^ providing empirical support for the use of Ru(II) complexes as potential FINs. Building on this insight, and inspired by the design of boron-containing liposomes (boronsomes), for improved intracellular delivery of boron molecules,^21, 22^ we designed a unique polypyridyl Ru(II)-based metallosurffactant^23, 24^ [Ru(bipy)_2_(*N*^4^,*N*^4’^-bis(2-oleamidoethyl)-2,2’-bipyridine-4,4’-dicarboxamide)](PF_6_)_2_ (**RuOA2**, Scheme S1). This complex self-assembles into liposome-like structures, significantly enhancing cellular delivery and driving potent ferroptosis activation through ROS generation in cancer cells (Figure 1).

As presented in Figure 1, the positively charged [Ru(bipy)_3_]^2+^ (Ru-trisBP) unit serves as the headgroup of the metallosurfactant. Unlike conventional designs that employ alkyl chains as tails, **RuOA2** incorporates two oleate chains derived from oleic acid (OA)−an abundant component in cancer cell plasma membranes, thus enhancing cancer cell targeting.^25^ The optimized structure of **RuOA2** (Figure 2a) features an octahedral headgroup with a diameter of 12.5 Å and the flexible tails measuring 23.2 Å facilitating conformational adaptability. This large headgroup extends the overall length of **RuOA2** to 35.8 Å, considerably longer than typical phospholipid designs. Molecular dynamics (MD) simulations reveal that **RuOA2** self-assemblies into a lipid-bilayer-like structure in aqueous solutions, with some OA tails oriented outward (Figure 2b). The bilayer’s thickness is approximately 78.25 Å–nearly double that of a traditional phospholipid bilayer. The spacing between adjacent headgroups (Figure 2c) indicates a densely packed assembly. Experimentally, UV-Vis spectra of **RuOA2** show hypochromicity and a redshift in the π-π* (∼280 nm) transitions of the bipyridine ligands, along with metal-to-ligand charge transfer transitions (∼480 nm) in 1XPBS compared to in methanol (Figure S1), suggesting extended bipyridine J-aggregate stacking, consistent with the simulated tight packing of [Ru(bipy)_3_]²⁺ units along the bilayer surfaces. Upon excitation at 450 nm, **RuOA2** emits bright red phosphorescence in water and shows no sensitivity to oxygen quenching (Figure S2), further indicating that the self-assembled close packing effectively shields the excited states from oxygen.

**Figure 2.**
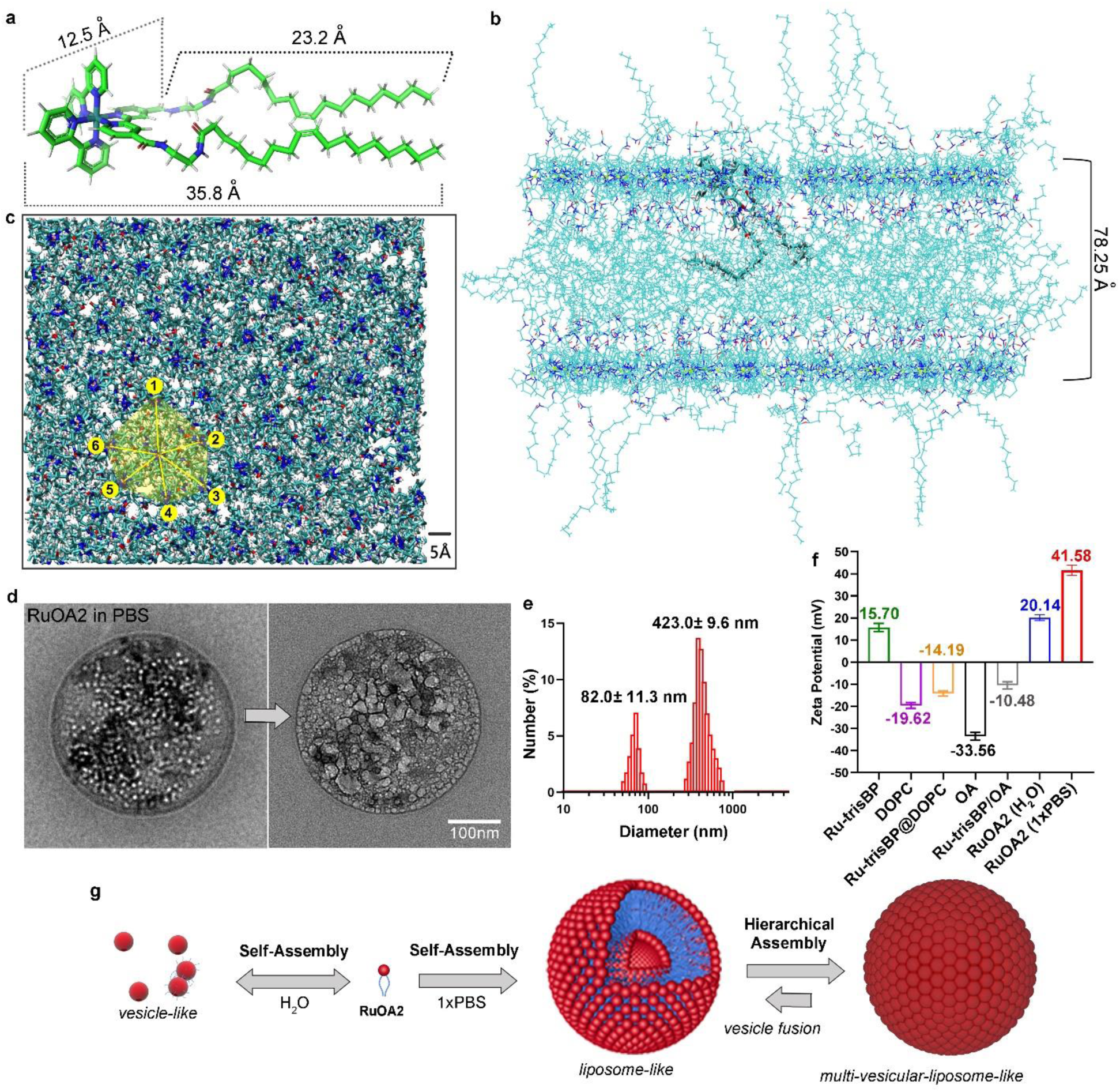
Dynamic self-assembly of RuOA2 in aqueous environment. **(a)** Spatial configuration and structural parameters of RuOA2. Cross-section **(b)** and top view **(c)** representations of bilayers formed from the self-assembly of RuOA2, based on a 128-monomer MD simulation in 3200 water molecules. In panel **b**, most RuOA2 molecules are depicted in wireframe mode, with one molecule shown in stick mode. In panel **c**, intermolecular distances between Ru centers are highlighted in yellow, with number labels corresponding to distances 12.31, 9.81, 12.62, 9.87, 9.94, 11.25 Å, respectively. **(d)** TEM images of the ruthenosome in PBS under weak electron beam and after electron beam irradiation for a few minutes. **(e)** Diameter distribution of RuOA2 self-assemblies in 1XPBS buffer as determined by DLS analysis. **(f)** Zeta potentials of Ru-trisBP, DOPC, OA, Ru-trisBP@DOPC, Ru-trisBP@OA and RuOA2 in 1XPBS buffer at a concentration of 20 μM, alongside RuOA2 in water at the same concentration. Mean ± SD (n=3). **(g)** Schematic depiction of the dynamic self-assembly process of RuOA2 in aqueous environment.

Experimentally, **RuOA2** self-assembled into particle clusters with an average size of ∼42.0 nm in water (Figure S3a, S3b). After negative staining with uranyl acetate (UAc), these self-assemblies transformed into evenly distributed vesicles with an average size of ∼32.0 nm, indicating that ionic conditions significantly influence the assembly process. In 1XPBS, which simulates physiological conditions, **RuOA2** self-assembled into liposome-like vesicles (Figure 2d and S3c) with a predominant diameter of ∼423 nm (Figure 2e). This size is significantly larger than liposomes formed from 1,2-dioleoyl-*sn*-glycero-3-phosphocholine (DOPC) and OA encapsulating Ru-trisBP, which exhibit average diameters of ∼335 nm and ∼90 nm, respectively (Figure S3a and S3b). Similar to these liposomes, **RuOA2** self-assemblies adopt a bilayer structure. However, unlike the typically negative surface charges seen in Ru-trisBP@DOPC and Ru-trisBP@OA liposomes, zeta potential measurements reveal that **RuOA2**-assembled ruthenosomes possess a high positive charge (+41.58 mV) (Figure 2f), which offers a significant advantage for mitochondrial targeting.^26^

During TEM imaging, we observed that **RuOA2**-assembled structure exhibited dynamic instability under electron beam irradiation. Exposure to the electron beam triggered a fission event within the ruthenosomes, leading to the formation of multiple smaller vesicles inside the original structure. Remarkably, these newly formed vesicles did not disperse, likely due to strong hydrophobic interactions between the outward oleate chains on their surfaces. These interactions caused the smaller vesicles to adhere to one another, maintaining the integrity of the ruthenosome as a multi-vesicular, liposome-like structure (Figures 2d, S3c and S3d). With continued irradiation, a fusion process was observed, where the small vesicles began to merge, gradually coalescing back into larger liposome-like structures (Video S1). This behavior highlights the dynamic nature of **RuOA2** self-assemblies, capable of undergoing transitions between fission and fusion, suggesting that the self-assembly of **RuOA2** is highly responsive to environmental conditions (Figure 2g).

### RuOA2 ruthenosome enhanced cellular uptake and mitochondrial targeting

**RuOA2** ruthenosomes exhibited broad cytotoxicity against a range of cancer cell lines commonly used as *in vitro* tumor models, including HeLa (cervical cancer), HepG2 (liver cancer), MCF7 (breast cancer), as well as cell lines from refractory cancers such as hypermutated CT26 and HCT116 (colorectal cancer, CRC), H1975 and A549 (lung cancer), and drug-resistant PANC-1 (pancreatic cancer). Notably, the IC50 values for **RuOA2** were below 20 μM for all these cancer cells, with the greatest potency observed in CRC cells, achieving sub-micromolar IC50 values (0.33±0.008 μM) (Figure S4). CRC is the third leading cause of cancer-related deaths worldwide, characterized by a high frequency of mutations and significant heterogeneity in immune responses and metabolic profiles among patients. Given the pressing need to unravel the mechanisms regulating ferroptosis in CRC, as well as to harness these pathways for drug development and potential clinical translation,^27^ we focused further investigation on HCT116 cells to explore **RuOA2** ruthenosome’s role in ferroptosis regulation.

Upon introducing **RuOA2** to HCT116 cells, we observed rapid accumulation of **RuOA2** at the plasma membrane within 10-20 minutes (Figure 3a). Closer examination via fluorescence imaging revealed transmembrane localization of **RuOA2** after 30 minutes of treatment (Figure 3b). By 1 hour, **RuOA2** remained membrane-bound, but a significant portion had internalized into the cell, with some colocalizing with mitochondria (Figure 3c and 3d). After an additional hour, the majority of **RuOA2** was fully internalized and predominantly localized within mitochondria. By the 4-hour mark, no **RuOA2** was detected at the plasma membrane, and all had accumulated in mitochondria, which had assumed a blob-like morphology instead of their typical tubular structure, indicating mitochondrial stress likely due to elevated mitochondrial ROS (mtROS) levels^28^. In comparison, Ru-trisBP@DOPC showed higher cellular delivery than Ru-trisBP@OA, but exhibited nonspecific localization in both mitochondria and lysosomes (Figure S5).

**Figure 3.**
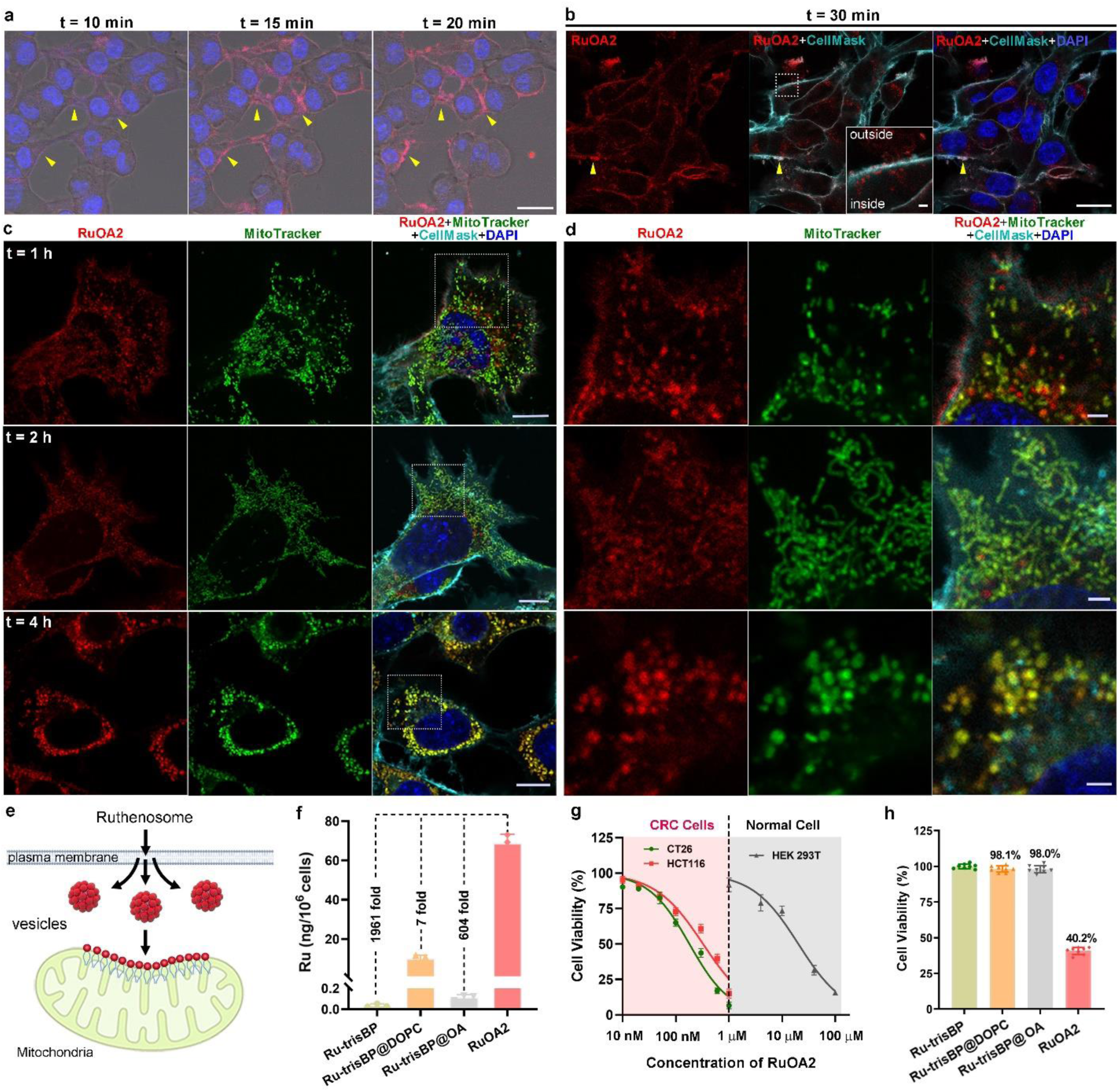
Intracellular delivery of RuOA2 to mitochondria in vitro. **(a)** Merged bright field and confocal fluorescence images of HCT116 cells upon the treatment of 5 μM RuOA2 (red) for 10, 15, and 20 minutes, co-stained with Hoechst (blue). The scale bar represents 20 μm. **(b)** Confocal fluorescence images of HCT116 cells upon the treatment of 5 μM RuOA2 (red) for 30 minutes, co-stained with Cell Mask (cyan) and DAPI (blue). The scale bars represent 10 μm. **(c)** Confocal fluorescence images of HCT116 cells treated with 5 μM RuOA2 (red) for 1, 2, and 4 hours, co-stained with MitoTracker (green), CellMask (cyan), and DAPI (blue). Scale bars represent 10 μm. **(d)** Zoomed-in images of white dash line squares in panel (c). Scale bars represent 2 μm. **(e)** Schematic depiction of intracellular delivery of RuOA2 via transition of ruthenosome into vesicles targeting mitochondria. **(f)** The cellular uptake of Ru in HCT116 cells upon the treatment of Ru-trisBP, Ru-trisBP@DOPC, Ru-trisBP@OA, and RuOA2 at a concentration of 1 μM for 4 hours through ICP-MS analysis. Mean ± SD (n=3). **(g)** HCT116, CT26 and HEK 293T cell viabilities upon the treatment of RuOA2 at various concentrations for 24 h. Mean ± SD (n=8). **(h)** HCT116 cell viability upon the treatment of Ru-trisBP, Ru-trisBP@DOPC, Ru-trisBP@OA, and RuOA2 at a concentration of 400 nM for 24 hours. Mean ± SD (n=8).

As summarized in Figure 3e, **RuOA2** enters the cell through a membrane-fusion-like process, followed by the release of **RuOA2** assemblies into the cytoplasma. These assemblies, driven by electrostatic attraction, accumulate at the mitochondria, triggering stress-induced morphological changes. The efficiency of cellular delivery was quantified via ICP-MS analysis (Figure 3f), showing that ruthenosomes enhanced the uptake of Ru-trisBP by nearly 2000-fold and increased uptake 6 times compared to Ru-trisBP@DOPC liposomes. The significant improvement in Ru-trisBP cellular delivery by ruthenosomes, more than 600-fold compared to Ru-trisBP@OA, highlights the advantages of **RuOA2**’s molecular design. Moreover, the high selectivity for mitochondrial targeting emphasizes the precision and effectiveness of this delivery system. In contrast, traditional liposome-mediated delivery of Ru-trisBP failed to induce cytotoxicity in HCT116 cells (Figure 3h). Interestingly, **RuOA2** exhibited potent cytotoxicity against CRC cell lines CT26 and HCT116 at nanomolar concentrations while sparing normal cells (HEK 293T), demonstrating strong cancer cell selectivity (Figure 3g).

### Mitochondria-targeted ruthenosomes induce ferritinophagy-mediated cancer suppression

To access whether **RuOA2** ruthenosomes effectively induce ferropotosis via mitochondrial targeting, we evaluated two hallmarks: increased levels of labile ferrous ion (Fe^2+^) levels and lipid peroxidation. Intracellular Fe^2+^ levels were measured using FeRhoNox-1, a non-fluorescent probe that specifically reacts with Fe^2+^ to emit fluorescence. Compared to untreated HCT116 cells and those treated with Ru-trisBP@DOPC or Ru-trisBP@OA, which showed minimal fluorescence, **RuOA2**-treated cells exhibited a marked increase in fluorescence, indicating elevated Fe^2+^ levels (Figure 4a and Figure S6). Lipid peroxidation was detected using the C11-BODIPY probe,^29^ where **RuOA2** treatment caused a fluorescence shift from red to green, indicating lipid peroxidation (Figure 4b and Figure S7). To further investigate **RuOA2**-induced ferroptosis, RNA sequencing (RNA-seq) was performed on HCT116 cells treated with 300 nM **RuOA2** for 24 hours. The analysis revealed insights into multiple cellular processes, including ferroptosis, endocytosis, and autophagy (Figure S8).

**Figure 4.**
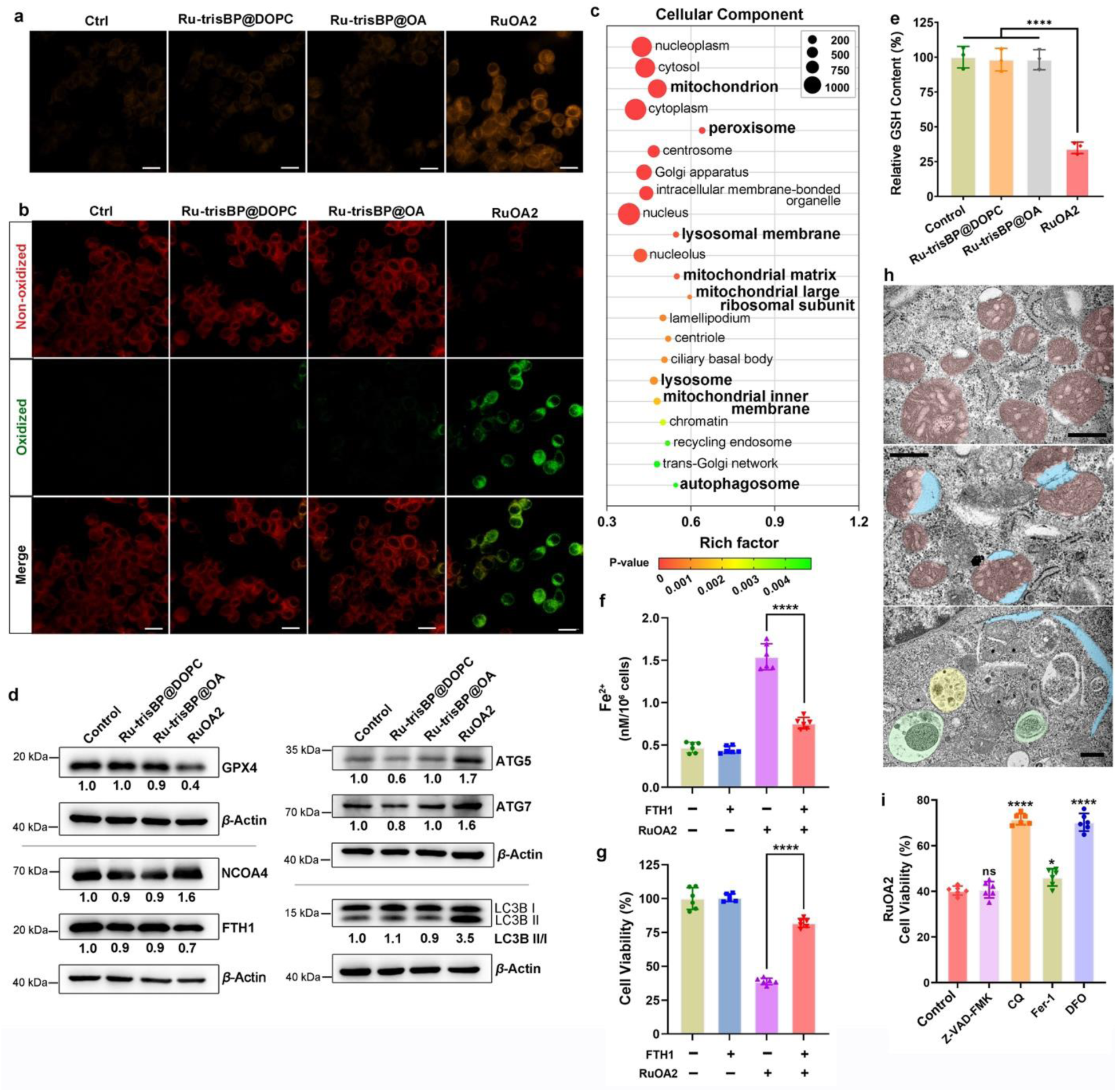
RuOA2 simulates ferritinophagy-mediated cancer suppression. **(a)** Fluorescent images of HCT116 cells with and without the treatments of Ru-complexes (300 nM, 24 hours) stained with FeRhoNox-1. Scale bars represent 20 μm. **(b)** Confocal fluorescence images of HCT116 cells incubated with C11-BODIPY after treatment with Ru-complexes at a concentration of 300 nM for 24 hours. The fluorescence transition from red to green indicated significant mitochondrial damage. Scale bars represent 20 μm. **(c)** Cellular component (CC) of GO enrichment analysis of differentially expressed genes in HCT116 cells treated with RuOA2 (300 nM) for 24 hours compared with the control. Bubble diagram was made based on the genes shown the top 30 CCs with high gene number. Bubble size represents the number of genes, while color represents the rich factor of gene expression. **(d)** Immunoblots for GPX4, NCOA4, FTH1, ATG5, ATG7, and LC3B I/II expression in HCT116 cells upon the treatment of Ru-complexes at a concentration of 300 nM for 24 hours. *β*-actin serves as the loading control. Immunoblots repeated in triplicate with similar results. **(e)** Relative intracellular GSH level in HCT116 cells upon treatment with Ru-trisBP@DOPC, Ru-trisBP@OA, and RuOA2 at a concentration of 300 nM for 24 hours. Mean ± SD (n=3). **(f)** The ferrous ion content in HCT116 cells transfected with control vector or FTH1-overexpression encoding plasmids with or without the treatment of 300 nM RuOA2 for 24 hours. Mean ± SD (n=3). **(g)** Viability of HCT116 cells transfected with control vector or FTH1-overexpression encoding plasmids with or without the treatment of 300 nM RuOA2 for 24 hours. Mean ± SD (n=6). **(h)** TEM images (false color) of RuOA2 (1μM, 4 hours) treated HCT116 cells. The mitochondria are highlighted in pink, the phagophores are highlighted in light blue, the autophagosomes are highlighted in green, and the autolysosomes is highlighted in yellow. Scale bars represent 500 nm. **(i)** HCT116 cell viability upon the treatment of **RuOA2** at a concentration of 400 nM for 24 hours after the pre-treatment of different PCD inhibitors. Mean ± SD (n=8). ns: no significant, ***p < 0.001, **** p < 0.0001.

The Gene Ontology (GO) analysis of cellular components (CC) highlighted significant enrichment in mitochondria, peroxisome, lysosome, and autophagosome (Figure 4c), underscoring the critical roles of mitochondrial dysfunction and autophagic processes in the cellular response to **RuOA2**. Specifically, gene expression analysis revealed distinct trends in the regulation of redox homeostasis, iron metabolism, lipid metabolism, and autophagy (Figure S9), suggesting that **RuOA2** treatment promotes iron-dependent lipid peroxidation, autophagy, and mitochondrial dysfunction. Notably, **RuOA2** induced the downregulation of *GPX4*^6^ and *NQO1*^30^, two key regulators of redox balance. The reduction of *GPX4* is particularly important, as its loss leads to excessive ROS accumulation, a hallmark of ferroptotic cell death. Additionally, the downregulation of *CISD1* and *NFS1*,^31^ which regulate mitochondrial iron-sulfur cluster homeostasis, further supports the disruption of mitochondrial function and the induction of redox imbalance.

The upregulation of the *TFRC* gene^32^, which governs cellular iron uptake via receptor-mediated endocytosis, indicates enhanced iron acquisition, contributing to ferroptosis initiation. Furthermore, *ATF3*^33^ and *ACSL4*^34^−key genes involved in lipid metabolism and the execution of ferroptosis−were upregulated, while *SCD1*^35^, a gene that regulates lipid homeostasis, was downregulated, highlighting an increase in lipid peroxidation and greater sensitivity to ferroptotic cell death. RNA-seq analysis also revealed significant upregulation of autophagy-related genes such as *ATG5* and *ATG7*,^36^ indicating autophagic activation in **RuOA2**-treated cells. The upregulation of *HMOX1* further supports autophagy’s role in regulating cellular iron levels. Moreover, the downregulation of ferritin heavy chain 1 (*FTH1*)^37^, responsible for iron storage, combined with the upregulation of *NCOA4*,^38, 39^ a key mediator of ferritin degradation via autophagy, suggests that **RuOA2** induces ferritinophagy−a selective form of autophagy that releases stored iron. Collectively, these findings implicate ferritinophagy as a crucial mechanism in **RuOA2**-induced ferroptosis (Figure S10).

The expression levels of key regulatory proteins involved in ferritinophagy were analyzed via western blotting (Figure 4d). Compared to untreated HCT116 cells and those treated with Ru-trisBP@DOPC or Ru-trisBP@OA, **RuOA2** treatment led to a 60% reduction in GPX4 expression, indicating elevated ROS production. To directly assess ROS levels, the DCFH-A assay was conducted, revealing that **RuOA2** treatment significantly enhanced ROS production compared to both Ru-trisBP@DOPC and Ru-trisBP@OA (Figure S11 and S12). Additionally, intracellular levels of the antioxidant GSH were also measured, and **RuOA2** treatment resulted in a nearly 70% decrease in GSH levels, shifting the redox balance towards a more oxidative state (Figure 4e). To further evaluate mitochondrial dysfunction associated with oxidative stress, JC-1 staining was performed to access mitochondrial membrane potential (MMP).^26^ The data revealed a significant reduction in MMP following **RuOA2** treatment (Figure S13). In comparison with cisplatin (CDDP), an established anticancer drug known to induce ROS-mediated cancer cell death, **RuOA2** was markedly more effective at inducing mtROS (Figure S14), underscoring its potency as a robust ROS inducer capable of triggering ferroptosis.

Consistent with the RNA-seq analysis, treatment with **RuOA2** resulted in a 60% increase in NCOA4 and a 30% decrease in FTH1, along with a 70% increase in ATG5 and a 60% increase in ATG7 expression. Additionally, immunoblotting revealed the conversion of LC3B from its LC3B-I to LC3B-II form. The increase in LC3B-II serves as a reliable indicator of autophagosome formation,^40, 41^ and we observed a 3.5-fold increase in the LC3B-II/LC3B-I ratio following **RuOA2** treatment. Autophagosome formation was further confirmed using TEM imaging (Figure 4h).^42, 43^ **RuOA2** treatment induced notable swelling of mitochondria with rounded cristae (highlighted in pink), indicative of reduced MMP due to increased calcium ion influx. The process of autophagy, a conserved transport pathway, involves the sequestration of targeted structures through the formation of phagophores, which mature into autophagosomes and are subsequently delivered to lysosomes for degradation. The TEM images presented in Figure 4h show membrane expansion and sealing of phagophores (highlighted in blue), the formation of autophagosomes (highlighted in green), and the presence of autolysosomes (highlighted in yellow). Collectively, these findings suggest that **RuOA2** induces ferritinophagy in HCT116 cells.

To confirm that **RuOA2** induces cell death through the ferroptosis pathway, we investigated the effect of overexpressing FTH1 in HCT116 cells treated with **RuOA2** (Figure 4f, 4g and S15). In control cells, FTH1 overexpression had no significant impact on intracellular labile Fe^2+^ levels or cell viability. However, in **RuOA2**-treated cells, FTH1 overexpression reduced Fe^2+^ levels by nearly 50%, leading to a restoration of cell viability, thereby confirming the critical role of FTH1 in regulating **RuOA2**-induced ferroptosis. To further explore the involvement of autophagy in **RuOA2**-induced ferroptosis, we evaluated the effects of various cell death inhibitors on **RuOA2**-treated HCT116 cells. As shown in Figure 4i, cell viability was 40.33% following **RuOA2** treatment alone, remaining nearly unchanged with the pan-caspase and apoptosis inhibitor Z-VAD-FMK^44^ (40.66%) or the lipid hydroperoxide scavenger Fer-1 (46.11%). In contrast, treatment with the autophagy inhibitor CQ,^45^ which disrupts autophagosome-lysosome fusion, and the iron chelator DFO,^46^ which suppresses ROS accumulation to inhibit ferroptosis,^47^ significantly increased cell viability to 71.59% and 70.23%, respectively. These results confirm that ferritinophagy plays a critical role in **RuOA2**-mediated cancer cell suppression. Furthermore, we examined HeLa, HepG2, MCF7, H1975, A549 and CT26 cells (Figure S16), confirming that **RuOA2** acts as a FIN and broadly induces ferritonophagy across various cancer cell types.

### Ruthenosomes induce ferritinophagy-mediated CRC suppression *in vivo*

To access the *in vivo* efficacy of **RuOA2** ruthenosomes in suppressing CRC, a xenograft model was established by subcutaneously implanting CT26 cells into C57BL/6 mice. Prior to the xenograft study, we confirmed that **RuOA2** induced ferritinophagy-mediated cell death in CT26 cells (Figure S17-24), consistent with findings from HCT116 human colorectal cancer cells. Mice were divided into four treatment group: saline (control), Ru-trisBP@OA, CDDP, and **RuOA2**, following the designated injection regimen (Figure 5a). As shown in Figure 5b, tumors growth progressed steadily in the saline and Ru-trisBP@OA groups, while **RuOA2** treatment significantly inhibited tumor growth ourperforming CDDP (Figure 5b, S25-26). At the end of the 22-day study, tumors were excised and weighted (Figure 5c and S27), with tumor weights in the **RuOA2**-treated group significantly lower than in the other groups, indicating effective tumor suppression. Histological analysis via H&E staining revealed extensive tumor cell death and loss of tissue integrity in the **RuOA2** treatment group (Figure S28) demonstrating pronounced anticancer activity.

**Figure 5.**
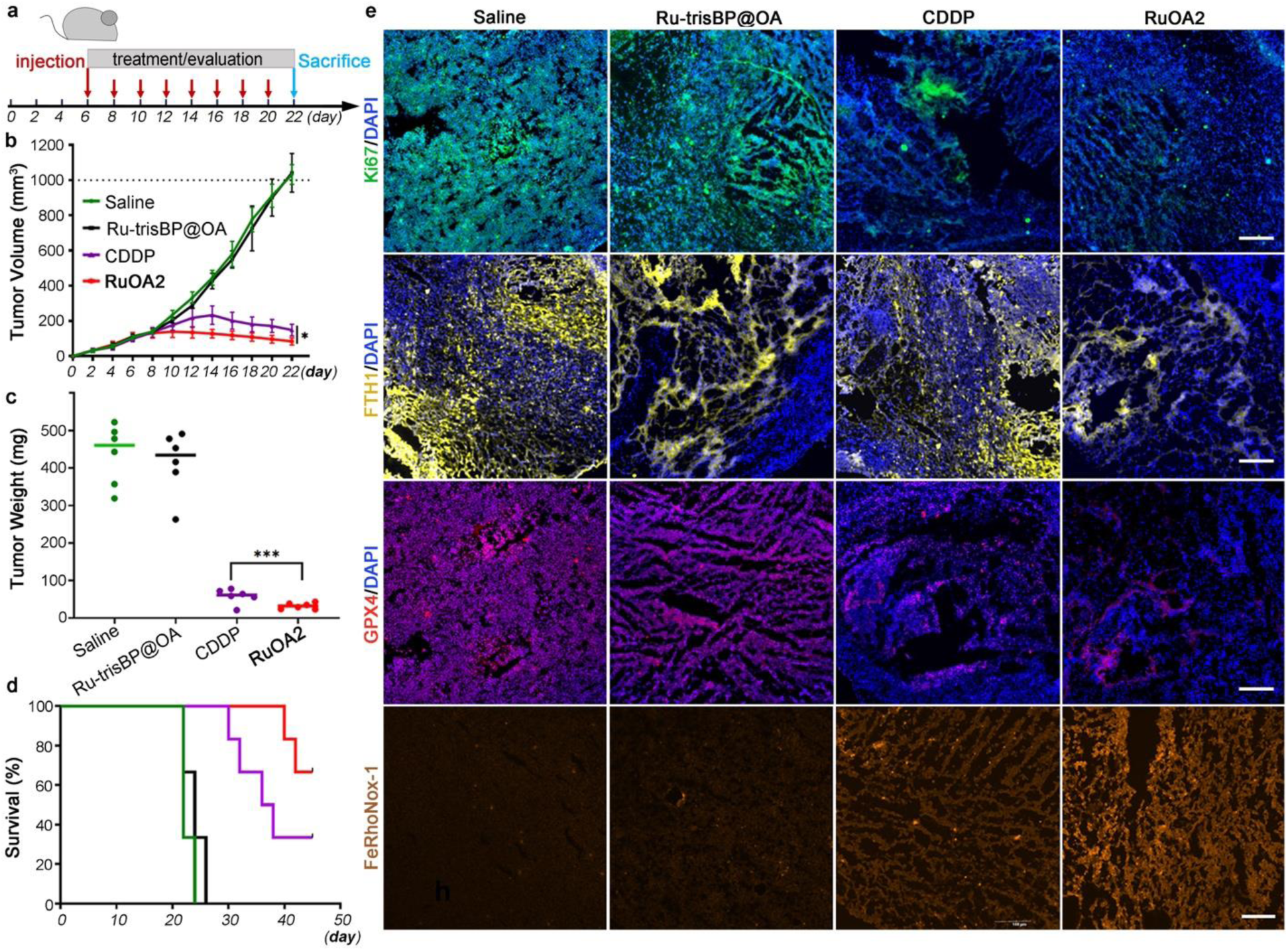
RuOA2 ruthenosome induces ferritinophagy-mediated CRC suppression *in vivo*. **(a)** Schematic representation of the injection regime in a CRC xenograft model using C57BL/6 mice. **(b)** Tumor growth was monitored via caliber measurements before each injection, with tumor volumes calculated and plotted. Mean ± SD (n=6). **(c)** Tumor weight of mice across different treatment groups at the end of 22 days. Mean ± SD (n=6). *p < 0.05, ***p < 0.001. **(d)** Survival rate statistics of mice under various treatments. (n=6). **(e)** Representative images of immunofluorescence staining for Ki67, FTH1, and GPX4, along with FeRhoNox-1 staining for detecting Fe^2+^ in tumor sections. Scale bars represent 100 μm.

Additionally, all treatment formulations had minimal impact on body weight, indicating good tolerability (Figure S29). Survival rates were monitored across treatment groups (Figure 5d), with the **RuOA2**-treated group exhibiting a markedly higher survival rate of 80% at 40 days, compared to 30% in the CDDP-treated group. Furthermore, analysis of serum biochemical markers (Figure S30) revealed that liver and kidney function remained within the healthy reference range, suggesting no detectable renal or hepatic toxicity in the **RuOA2**-treated mice. Post-mortem analysis of major organs (heart, liver, spleen, lungs, and kidneys) via H&E staining (Figure 31) revealed no signs of significant organ damage, further implying low systemic toxicity of **RuOA2**. Collectively, these results highlight the potent antitumor efficacy and favorable biosafety profile of RuOA2, demonstrating its potential as a therapeutic agent for CRC treatment.

To further investigate the molecular mechanisms underlying tumor suppression, we performed immunofluorescence staining and western blotting on tumor sections to assess the expression of the proliferation marker Ki67, the ferritin regulator FTH1, and the ferroptosis-related protein GPX4, alongside FeRhoNox-1 staining (Figure 5e, S32 and S33). The results revealed a significant reduction in Ki67 expression in the **RuOA2**-treated tumors, indicating decreased tumor cell proliferation. Additionally, FTH1 and GPX4 levels were notably downregulated, accompanied by increased FeRhoNox-1 fluorescence, reflecting elevated levels of labile Fe^2+^. These findings support the conclusion that **RuOA2** ruthenosomes effectively induce ferritinophagy-mediated tumor cell death *in vivo*, leading to more pronounced CRC suppression compared to CDDP.

## Discussion

The development of novel liposome systems has garnered significant attention due to their efficiency in delivering therapeutic agents into cells. However, many current designs, such as functional liposomes (e.g., boronsomes and liposome-in-liposome systems), lack specificity for subcellular organelles.^48^ In this study, we introduce a ruthenium-based liposomal system, termed ruthenosome, which represents the first ruthenium-based FIN that triggers ferroptotic tumor cell death without the need for light activation. Our design leverages the hierarchical self-assembly of **RuOA2** into multiscale vesicle-to-vesicle structures due to strong hydrophobic interactions between the outward-facing oleate chains on their surfaces. This unique molecular architecture not only facilitates enhanced cellular uptake but also achieves precise mitochondrial targeting, a critical advantage over conventional liposomal systems.

Ruthenosomes function as potent ROS generators, initiating ferroptosis and displaying broad applicability across various cancer types. The ability of **RuOA2** ruthenosomes to selectively target mitochondria and induce ferroptosis without external stimuli marks a significant breakthrough in metallodrug design. The promising in vivo results, including superior tumor suppression and minimal toxicity, underscore the therapeutic potential of RuOA2 in cancer treatment. This study highlights the versatility of ruthenosomes as multifunctional nanocarriers and paves the way for the development of the next generation of metallodrugs. By incorporating nanotechnology, future designs can further improve therapeutic efficacy while minimizing off-target effects, offering a transformative approach to targeted cancer therapies.

## Methods

### Synthesis of RuOA2

To a suspension of oleic acid (1 mL) in anhydrous acetonitrile (10 mL), **Ru-2** complex (103 mg, 0.1 mmol) was added, followed by HBTU (0.5 g) and DIC (0.2 mL). The reaction mixture was stirred and refluxed at 60 ℃ for 24 hours, then allowed to cool to room temperature. After removing the precipitates by filtration, the solvent was evaporated under reduced pressure obtaining solid residue, which was resuspended in 10 mL of anhydrous acetonitrile. Any remaining insoluble material was filtered off. The crude product was purified by column chromatography on aluminum oxide to yield pure **RuOA2** (92.2 mg) as a dark brown powder with a 59.0% yield. Details on the synthesis of **Ru-2** are provided in the Supplementary Information

### Preparation of Ru-trisBP@DOPC and Ru-trisBP@OA liposomes

Ru-trisBP@DOPC liposomes were synthesized using a thin-film hydration method. Specifically, 27.64 mg of DOPC (D68710, Acmec, China) was dissolved in methanol, then evaporated at 40 ℃ for 30 minutes to form a uniform thin film. The dried lipid film was hydrated with 3 mL of a Ru-trisBP solution (0.5 mg/mL in water) and sonicated for 30 minutes. This mixture was then extruded through 0.45 μm pore-size polycarbonate membrane filters to produce Ru-trisBP@DOPC liposomes. Unencapsulated Ru-trisBP was removed by chromatography on a Sephadex G10 column. The final concentration of Ru-trisBP encapsulted in Ru-trisBP@DOPC liposomes was determined to be 1.98 mM, with a DOPC to Ru-trisBP molar ratio of 17.86:1, and a loading efficacy of 84%. Ru-trisBP@OA liposomes were prepared by mixing Ru-trisBP and oleic acid (OA) at a 1:2 molar ratio in deionized water to mimic the Ru-trisBP/OA ratio found in the structure of **RuOA2**. The mixture was then subjected to continuous sonication for 30 minutes, facilitating co-assembly and yielding Ru-trisBP@OA liposomes.

### Preparation of RuOA2 ruthenosomes

**RuOA2** was first dissolved in dimethyl sulfoxide (DMSO) and then diluted at a 1:1000 ratio with deionized water or 1XPBS. The resulting solution was sonicated for 30 minutes to obtain ruthenosomes.

### Simulation

The conformational optimization of **RuOA2** was conducted using ORCA software with density functional theory (DFT), employing the B3LYP functional and the def2-SVP basis set. Molecular dynamics simulations were then carried out to model the self-assembly of **RuOA2**. A supercell was constructed with 8 unit cells along both the x and y directions. Each unit cell contained two **RuOA2** molecules arranged in an up-and-down mirror plane orientation, with the Ru atoms end facing outward. The initial dimensions of the supercell were 144 Å x 160 Å x 46 Å in the x, y, and z directions, respectively. Due to the limited availability of force field parameters for PF_6_^−^ ions, Cl^−^ ions were used as substitutes. The supercell was filled with 3200 water molecules and 256 Cl^-^ ions using Packmol software. Molecular dynamics simulations were performed using LAMMPS software with the PVFF force field, applying Lennard-Jones potential with a cutoff distance of 10 Å. Bond, angle, dihedral, and improper interactions were defined using class2 force field types. The NVT ensembles was employed with a temperature of 300 K, allowing the simulation box to adjust its size dynamically. Over the course of the simulation, the box size decreased, ultimately reaching final dimensions of 70 Å x 80 Å x 47 Å in the x, y, and z directions, respectively.

### TEM imaging of liposomes and ruthenosomes

Following the preparation protocols detailed above, Ru-trisBP@DOPC liposomes (10 μM Ru-trisBP), Ru-trisBP@OA (10 μM), and **RuOA2** ruthenosomes (10 μM) were prepared. After allowing the samples to stabilize at room temperature for 2 hours, 10 μL aliquots of sample solution were deposited onto glow-discharge copper grids (400 mesh) coated with a thin carbon film. The samples were left on the grid for 30 seconds before excess solution was removed. The grids were then rinsed with deionized water up to three times. TEM images were acquired under high vacuum using a Talos-L120C transmission electron microscope (Thermo Fisher). The dynamic morphological transitions of ruthenosomes were captured with a multipurpose electron microscope (JEM-F200, JEOL, Japan).

### Cell culture

HCT116, CT26, and PANC-1 were obtained from the Cell Bank of the Chinese Academy of Sciences. HeLa, HepG2, MCF7, H1975 and A549 were generously provided by the Huadong Liu’s lab at Xi’an Jiaotong University. PANC-1, HeLa, HepG2, MCF7, and A549 cells were cultured in Dulbecco’s Modified Eagle’s Medium (DMEM), while HCT116 cells were maintained in McCoy’s 5a medium. H1975 and CT26 cells were cultured in RPMI 1640 medium. All culture media were supplemented with 10% fetal bovine serum (FBS) and 1% penicillin/streptomycin. Cells were incubated at 37 °C in a humidified atmosphere with 5% CO₂ and passaged every 3 days using a 0.25% trypsin-EDTA solution.

### Cell viability assay

Cells in the exponential growth phase were seeded into a 96-well cell culture plate at a density of 5×10^3^ cells per well and incubated for 24 hours at 37 °C in 5% CO_2_ atmosphere. Molecules, liposomes and ruthenosomes were added to the wells at the specific concentrations. After the desired incubation period, 10 μL of MTT solution (5 mg/mL) was added in each well. The plates were then incubated at 37 °C for 4 hours. Subsequently, the MTT-containing supernatant was removed, and 100 μL of DMSO was added to dissolve the formazan crystals formed by viable cells. The resulting purple solution was measured for optical density at 490 nm using a SuPerMax 3100 plate reader (Shanpu, China).

### Intracellular uptake assessment

HCT116 cells were seeded in 6-well plates at a density of 5×10^4^ cells per well and incubated overnight. Ru-trisBP@DOPC (1 μM Ru-trisBP), Ru-trisBP@OA (1 μM), or **RuOA2** (1 μM) was added, and the cells were incubated at 37 ℃ for 4 hours. After incubation, cells were washed with PBS, and then harvested, lyophilized and subsequently subjected to microwave digestion. The resulting residue was reconstituted in deionized water to a final volume of 10 mL, generating the test solution. A specified volume of this solution was then analyzed for Ru content using the Inductively Coupled Plasma Mass Spectrometry (ICP-MS) (PerkinElmer NexION 300X).

### RNA sequencing and bioinformatics analysis

HCT116 cells were seeded in 6-well plates at a density of 5×10^4^ cells per well and treated with 300 nM **RuOA2** for 24 hours. Following treatment, cells were harvested, and total RNA was extracted using TRNzol Universal Reagent (DP424, Tiangen, China). RNA-seq was performed by Lc-Bio Technologies. Genes with false discovery rates (FDR) < 0.05 and lengths > 200 bp were considered to show differential expression. Kyoto Encyclopedia of Genes and Genomes (KEGG) and Gene Ontology (GO) enrichment analyses were conducted using the DAVID online database. Gene Set Enrichment Analysis (GSEA) followed the standard procedures outlined in the GSEA User Guide (http://www.broadinstitute.org/gsea/doc/GSEAUserGuideFrame.html).

### Intracellular glutathione (GSH) quantification assay

HCT116 cells were seeded in 6-well plates at a density of 5×10^4^ cells per well and incubation overnight. Cells were then treated with Ru-trisBP@DOPC (300 nM Ru-trisBP), Ru-trisBP@OA (300 nM) or **RuOA2** ruthenosome (300 nM) for 24 hours. The intracellular GSH levels were quantified using a GSH Assay Kit (A006-2-1, Njjcbio, China). For the calibration curve, 1 mM GSH standard solutions were diluted to prepare solutions at various concentrations (0, 5,10, 20, 50, and 100 μM). After treatment, cells were harvested, and colorimetric quantitative of GSH was performed at 405 nm.

### Confocal microscopy and intracellular localization analysis

Cells in the exponential growth phase were seeded into 35 mm glass bottom dishes at a density of 2×10^4^ cells per dish. Once fully adhered, the culture medium was replaced with fresh medium containing various concentrations of liposomes, ruthenosomes, or molecules. After the desired incubation time, cells were washed three times with PBS and subsequently stained with cell labeling dyes according to the manufacturer’s protocols. Fluorescent images were acquired using an Olympus FV3000 confocal laser scanning microscope. For intracellular localization studies, the following stains were used:

Nuclear staining: Hoechst 33258 (1 µg/mL; H4047, UElandy, China) or DAPI (1 µg/mL; C1002, Beyotime, China) with excitation/emission at 405/460 ± 20 nm.

Mitochondrial staining: MitoTracker Green (50 nM; A66441, Thermo Fisher Scientific, USA) with excitation/emission at 488/516 ± 10 nm.

Cell Membrane staining: CellMask™ Deep Red stain (50 nM; A57245, Thermo Fisher Scientific) with excitation/emission at 640/670± 20 nm.

Lysosome staining: Lyso Tracker Green (50 nM, L7526, Thermo Fisher Scientific) with excitation/emission at 488/511±10 nm.

### Mitochondrial membrane potential assessment

The mitochondrial membrane potential was measured using the JC-1 dye (C2006, Beyotime) following the manufacturer’s protocol. Briefly, cells were incubated with a 1×JC-1 working solution at 37 °C for 30 minutes. Fluorescence settings were configured with excitation/emission at 488/535 ± 20 nm for JC-1 monomers and 488/570 ± 20 nm for JC-1 aggregates, to differentiate between polarized and depolarized mitochondrial states.

### ROS, Lipid persoxidation (LPO), and Fe^2+^ quantification assay

To detect reactive oxygen species (ROS), 10 µM DCFH-DA (S0033S, Beyotime) was used with fluorescence settings of Ex/Em 488/525 ± 20 nm. For lipid peroxidation (LPO) detection, 5 μM C11 BODIPY (D3861, Thermo Fisher Scientific) was employed, with Ex/Em 561/591 ± 10 nm (red channel) and Ex/Em 488/510 ± 10 nm (green channel). Fe^2+^ levels were assessed using 5 μM FeRhoNox-1 (MX4558, MKBio, China) with Ex/Em settings at 561/580 ± 10 nm. Images were processed and analyzed with ImageJ software. For quantitative analysis, background subtraction and normalization were applied, and the integrated density of fluorescence signals was measured using ImageJ.

### TEM imaging of cells

HCT116 cells were cultured in 10 cm dishes for 24 hours and treated with 300 nM **RuOA2** for an additional 24 hours. Cells were harvested by trypsinization and fixed in 2.5% glutaraldehyde, followed by post-fixation with 1.0 % osmium tetroxide. Samples were then embedded in epoxy resin and sectioned into 70 nm slices. TEM images were captured using a Hitachi H-7650 TEM at an acceleration voltage of 80 kV.

### Overexpression of FTH1 in HCT116 and CT26 cells

HCT116 and CT26 cells were seeded in 6-well plates and allowed to grow to 70-80% confluence. The FTH1-GFP expression plasmid was synthesized by Sangon Biotech. For each well, 2 µg of plasmid DNA was diluted in 125 µL of Opti-MEM I Reduced Serum Medium (Gibco). Separately, 5 µL of Lipo 6000™ Transfection Reagent (C0526FT, Beyotime, China) was diluted in 125 µL of Opti-MEM I. After a 5-minute incubation at room temperature, the diluted DNA and Lipo 6000™ were mixed and incubated for 20 minutes to allow the formation of DNA-lipid complexes. The mixture was then added dropwise to each well containing cells in serum-free medium and incubated for 6 hours at 37°C in a CO₂ incubator. Following this, the transfection medium was replaced with fresh complete growth medium, and the cells were incubated for an additional 24-48 hours prior to downstream analysis.

### Western blot analysis

Cells and tumor tissues were lysed in cell lysis buffer with protease inhibitors (P0013, Beyotime), followed by centrifugation at 12000 rpm for 10 minutes at 4 ℃ . The supernatant was collected, and protein concentrations were determined using a BCA Quantitation Kit (23228, Thermo Fisher Scientific). Proteins were then mixed with SDS sample buffer containing 10% β-mercaptoethanol, separated by SDS-PAGE, and transferred to nitrocellulose membranes for immunoblotting. β-Actin was used as the loading control. Primary antibodies included: anti-β-Actin (66009-1-Ig, Proteintech, 1:5000), anti-LC3B I/II (ET1701-65, HuaBio, 1:1000), anti-ATG5 (ET1611-38, HuaBio, 1:1000), anti-ATG7 (ET1610-53, HuaBio, 1:1000), anti-NCOA4 (ER62707, HuaBio, 1:2000); anti-FTH1 (T55648, Abmart, 1:1000); anti-GPX4 (T56959S, Abmart, 1:1000). Blots were developed using an enhanced chemiluminescence (ECL) detection system (1705061, Bio-Rad, USA), and signal were visualized on a Chemiluminescence imaging analysis system (BG-gdsAUTO 720, Baygene, China). Band intensities were quantified using ImageJ software.

### Immunohistochemistry

Tumor tissues were collected and embedded in Optimal Cutting Temperature (OCT) compound. OCT-embedded tissues were sectioned at a thickness of 5 μm and mounted onto glass slides. Sections were fixed in 4% paraformaldehyde for 20 minutes, permeabilized with 0.1% Triton X-100 for 1 hour, and blocked with 5% bovine serum albumin (BSA) at room temperature for 30 minutes. The sections were then incubated overnight at 4 °C with primary antibodies against FTH1 (1:500), GPX4 (1:500), and Ki67 (PT0321R, Immunoway, 1:500), followed by incubation with a Cy3-conjugated goat anti-rabbit IgG secondary antibody (SA00009-2, Proteintech, 1:100) for 1 hour at room temperature. Nuclear staining was performed using 4′,6-diamidino-2-phenylindole (DAPI, P0131, Beyotime) for 30 minutes. Confocal images were captured on an Olympus FV3000 laser scanning microscope.

### Animal studies

All animal experiments were approved by the Animal Ethics Committee of Xi’an Jiaotong University (No. 2021-1123). Female C57BL/6 mice (6-8 weeks) were obtained from the Experimental Animal Center of Xi’an Jiaotong University and housed under specific pathogen-free conditions with a 12-hour light/dark cycle and free access to food and water. Six days after tumor cell inoculation, when tumor signals became detectable by *in vivo* imaging, mice were randomized into four groups and peritumorally injected every other day with a 5 mg/kg dose of liposome, ruthenosomes, or drug molecules. On day 22, two days after the last treatment, tumor tissues were collected for hematoxylin and eosin (H&E) staining to evaluate antitumor effects.

### Xenografts implantation

To generate murine subcutaneous tumors, 5×10^6^ CT26 cells stably expressing the GFP-Luc fusion gene (CT26-luciferase cells) were injected subcutaneously into female C57BL/6 mice aged 8 weeks.

### In vivo bioluminescence imaging

On day 6 post-inoculation, detectable tumor signals were imaged using the *in vivo* imaging system. Mice were divided into four groups (Saline, Ru-trisBP@OA, CDDP, and RuOA2). Bioluminescence images were captured on days 6, 14, and 22 following an intraperitoneal injection of D-Luciferin potassium salt (150 mg/kg) using the VISQUE In VIVO Smart-LF imaging system.

### Therapeutic evaluation

Tumor growth was monitored by measuring tumor weight, length (L), and width (W). Tumor volume was calculated with the formula: 1/2(L × W^2^). Mice body weights were recorded every 2 days to monitor health. At the end of the treatment period, mice were euthanized, and tumor tissues were stained with H&E for morphological evaluation. Survival time was defined as the duration from tumor implantation until sacrifice or when tumor volume reached 1000 mm³, with a maximum follow-up of 45 days.

### Statistics and reproducibility

Data were analyzed using GraphPad Prism (version 9.0) and are presented as the mean ± standard deviation (SD) with n ≥ 3 unless specified otherwise. Unpaired Student’s t-tests were used for comparisons the between two groups, and one-way analysis of variance (ANOVA) was used for multiple group. A p-value of less than 0.05 was considered statistically significant. Statistical significance is indicated by asterisks: p < 0.05, **p < 0.01, ***p < 0.001, ****p < 0.0001.

## Author Contributions

M. Z., Y. Z. and G. L. conceived the study and designed the experiments. C. M., S. L. and Y. M., performed the experiments. C. M., S.L. and H. Y. conducted Spectral analysis, cell studies, and animal studies, L. S. and B. Z. conducted molecular simulation experiments, J. S. and J. Z. conducted TEM imaging, L. Y. conducted immunofluorescence assay, X. L. conducted confocal microscopy. M. Z. provided constructive discussion on clinical colorectal cancer treatment. Y. Z., C. M. and G. L. wrote and revised the paper. All authors have approved the final version of the manuscript.

## Acknowledgement

This work was supported by financial support from the open research fund of Songshan Lake Materials Laboratory (2022SLABFK05), the National Natural Science Foundation of China (No. 22107087). We thank the Chao Li at the Instrument Analysis Center of Xi’ an Jiaotong University for the assistance in TEM imaging experiments, and Baochang Lai at the Cardiovascular Research Center of Xi’an Jiaotong University for the assistance in confocal microscope imaging experiments.

## Supplemental Information

Supplemental information includes Supplemental Experimental Procedures, Supplementary Tables, and Supplementary Figures can be found with this article online at: XXX.

