## Supplemental files for "Assembling Ruthenium Complexes to Form Ruthenosome Unleashing Ferritinophagy-mediated Tumor Suppression"

The supplementary information contains: 1 scheme, 33 figures and 8 attachments.

### Materials and general instruments

All the solvents and chemicals are commercially available. Chemicals were used without further purification. $^{1}$H and $^{13}$C spectra were recorded in a deuterated solvent on Bruker AVANCE 400 MHz spectrometer or Bruker AVANCE III NEO 500 MHz spectrometer. $^{1}$H NMR chemical shift (δ) is given in ppm referring to internal standard tetramethylsilane (TMS). All coupling constants (*J*) are given in Hz. Electrospray ionization mass spectrometry (ESI-MS) was recorded on an LTQ XL linear ion trap mass spectrometer (Thermo Fisher). Electronic absorption spectra were recorded on an Agilent Cary 60 UV-Vis spectrophotometer. Steady-state emission spectra were recorded on a fluorescence spectrometer (Hamamatsu). Dynamic light scattering measurements were carried out on an Anton Paar Litesizer 500 dynamic light scattering (DLS) instrument equipped with a He-Ne laser (l=633nm) and a backscattering detector at a fixed angle of 173°. Cell cycle analysis was measured on a Beckman CytoFLEX-S flow cytometry.

### Synthesis

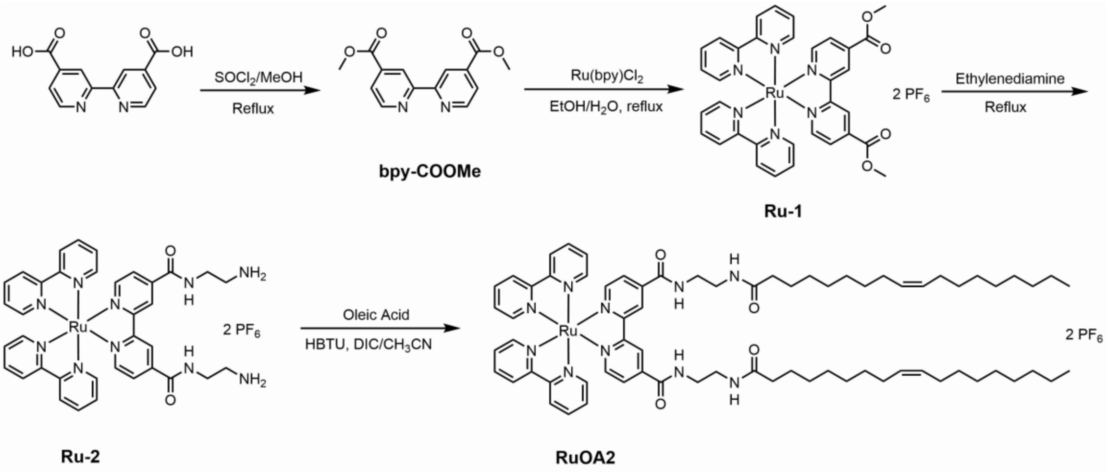


**Scheme S1.** Synthetic path of **RuOA2**.

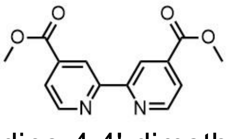

**bpy-COOMe**

**bpy-COOMe** (2,2'-bipyridine-4,4'-dimethylcarboxylate): 5 mL $SOCl_2$ was added to a suspension of 2,2'-bipyridine-4,4'-dicarboxylic acid (0.488 g, 2.0 mmol) and refluxed for 6hrs. Solvent was removed under reduced pressure to obtain a light-yellow solid. Yield 0.531 g, 97.5%.

$^1$H NMR (400 MHz, CDCl3) δ 9.05 (s, 2H), 8.92 (s, 2H), 7.98 (s, 2H), 4.02 (s, 6H). $^{13}$C NMR (101 MHz, $CDCl_3$) δ 165.07, 154.70, 149.58, 139.65, 123.98, 121.49, 53.05. (**Attachment 1**)

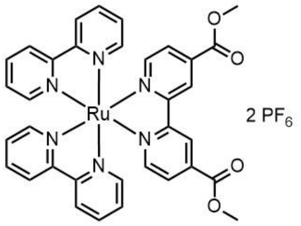


**Ru-1**

**Ru-1**: Mixture of bpy-COOMe (0.272 g, 1 mmol) and precursor cis-Ru(bpy)$_2$Cl$_2$ (0.484 g, 1 mmol) in EtOH/$H_2O$ (50 mL/10 mL) was refluxed at 90 ℃ overnight. Solvent was removed under reduced pressure and solid was dissolved in 10 mL water, to which $NH_4PF_6$ solution (0.5 g in 2 mL water) was added and stirred at r.t. for 10 min. Dark brown precipitate was filtered and washed with ice-water. Crude product was purified on neutral aluminum oxide (300 mesh) chromatography eluted with 1:2 acetonitrile/toluene. Yield 0.731 g, 75.0%.

ESI-MS calcd. for $C_{34}H_{28}F_{12}N_6O_4P_2Ru$ 975.63, found $[M-2PF_6]^{2+}$, m/z = 343.28. (**Attachment 2**) $^1$H NMR (400 MHz, $CD_3CN$) δ 9.03 (s, 2H), 8.50 (dd, *J* = 8.1, 4.4 Hz, 4H), 8.08 (tt, *J* = 7.8, 4.0 Hz, 4H), 7.94 (d, *J* = 5.8 Hz, 2H), 7.81 (dd, *J* = 5.8, 1.6 Hz, 2H), 7.67 (dd, *J* = 17.1, 5.4 Hz, 4H), 7.40 (dt, *J* = 19.8, 6.6 Hz, 4H), 3.99 (s, 6H). $^{13}$C NMR (101 MHz, $CD_3CN$) δ 164.53, 158.34, 157.36, 157.23, 153.50, 152.44, 152.18, 139.06, 138.90, 128.39, 128.30, 127.08, 125.03, 125.01, 124.29, 53.62. (**Attachment 3**)

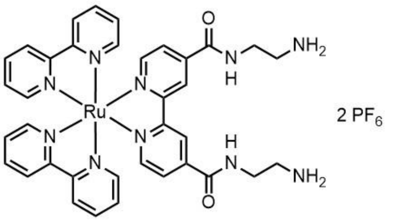


**Ru-2**

**Ru-2**: Complex Ru-1 (0.682 g, 0.7 mmol) was added to 10 mL dry ethylenediamine and refluxed at 110 ℃ under $N_2$ atmosphere for 12 hr. Solvent was removed under reduced pressure and 10mL dry acetonitrile was added. Insoluble salt precipitate was filtered off. Solvent was removed again under reduced pressure. Crude product was purified on neutral aluminum oxide (300 mesh) chromatography eluted with 2:1 acetonitrile/toluene. Yield 0.522 g, 72.2%.

ESI-MS calcd. for $C_{36}H_{36}F_{12}N_{10}O_2P_2Ru$ 1031.75, found $[M-2PF_6]^{2+}$, m/z = 371.45. (**Attachment 4**) $^1H$ NMR (400 MHz, $CD_3CN$) δ 9.89 (s, 2H), 8.53 (dd, *J* = 7.8, 5.5 Hz, 4H), 8.09 (dd, *J* = 17.0, 8.2 Hz, 4H), 7.85 (dd, *J* = 21.6, 5.8 Hz, 4H), 7.76 (t, *J* = 4.5 Hz, 4H), 7.43 (dt, *J* = 12.9, 6.5 Hz, 4H), 3.54 (t, *J* = 5.3 Hz, 4H), 2.99 (t, *J* = 5.6 Hz, 4H). $^{13}C$ NMR (101 MHz, $CD_3CN$) δ 163.76, 158.26, 157.43, 152.97, 152.45, 152.26, 142.65, 138.67, 128.28, 126.36, 124.92, 122.69, 42.01, 40.78. (**Attachment 5**)

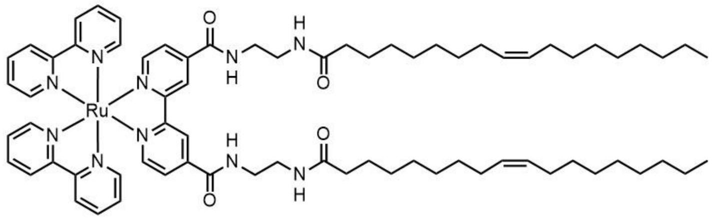


**RuOA2**

**RuOA2.** ESI-MS calcd. for $C_{72}H_{100}F_{12}N_{10}O_4P_2Ru$ 1560.65, found $[M-2PF_6]^{2+}$, m/z = 635.64, $[M-PF_6]^{+}$, m/z = 1415.64. (**Attachment 6**) $^1H$ NMR (400 MHz, MeOD) δ 9.11 (s, 2H), 8.73 (d, *J* = 6.3 Hz, 4H), 8.16 (dd, *J* = 13.5, 6.7 Hz, 4H), 8.00 (d, *J* = 5.9 Hz, 2H), 7.83 (d, *J* = 5.6 Hz, 6H), 7.58 – 7.44 (m, 4H), 5.41 – 5.29 (m, 4H), 3.51 (dt, *J* = 10.7, 5.1 Hz, 8H), 2.20 (t, *J* = 7.6 Hz, 4H), 2.03 (dd, *J* = 13.3, 6.8 Hz, 8H), 1.64 – 1.53 (m, 4H), 1.30 (d, *J* = 9.0 Hz, 40H), 0.91 (dd, *J* = 9.0, 4.5 Hz, 6H). $^{13}C$ NMR (101 MHz, MeOD) δ 175.66, 164.40, 157.56, 156.97, 156.90, 151.97, 151.37, 151.19, 142.84, 138.19, 129.51, 129.35, 127.73, 125.14, 124.35, 122.13, 40.23, 38.31, 35.80, 31.65, 29.43, 29.37, 29.34, 29.20, 29.06, 29.04, 28.94, 28.89, 28.81, 26.73, 25.55, 22.33, 13.05. (**Attachment 7**)

Counter ion $PF_6^-$ was confirmed by $^{19}F$ and $^{31}P$ NMR. $^{19}F$ NMR (376 MHz, MeOD) δ -73.49, -75.37. $^{31}P$ NMR (162 MHz, MeOD) δ -131.54, -135.90, -140.26, -144.63, -148.99, -153.35, -157.71. (**Attachment 8**)

### Supporting Figures

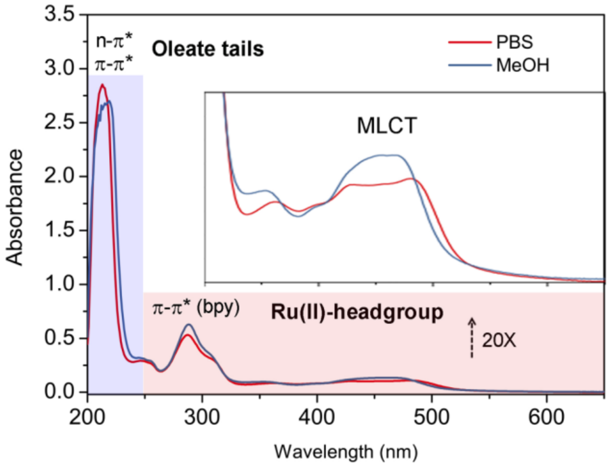


**Figure S1.** UV-vis absorption of RuOA2 dissolved in PBS or MeOH.

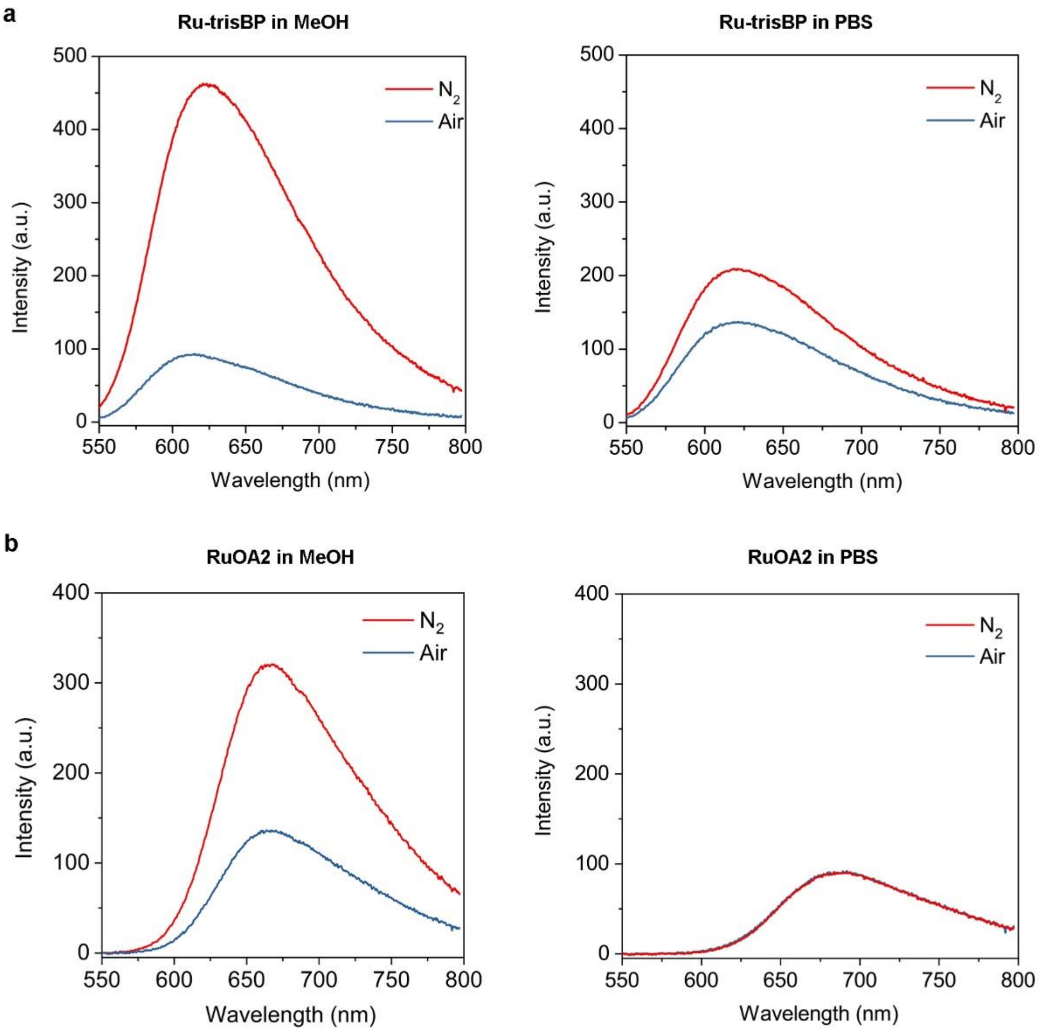


**Figure S2.** Emission spectra of 10 μM **Ru-trisBP** (a) or **RuOA2** (b) in MeOH with 0.1% DMSO or in PBS buffer with 0.1% DMSO. Solutions were bubbled with air (blue traces) or nitrogen (red traces) for 3 minutes prior to each measurement. Excitation wavelength was set to 450 nm.

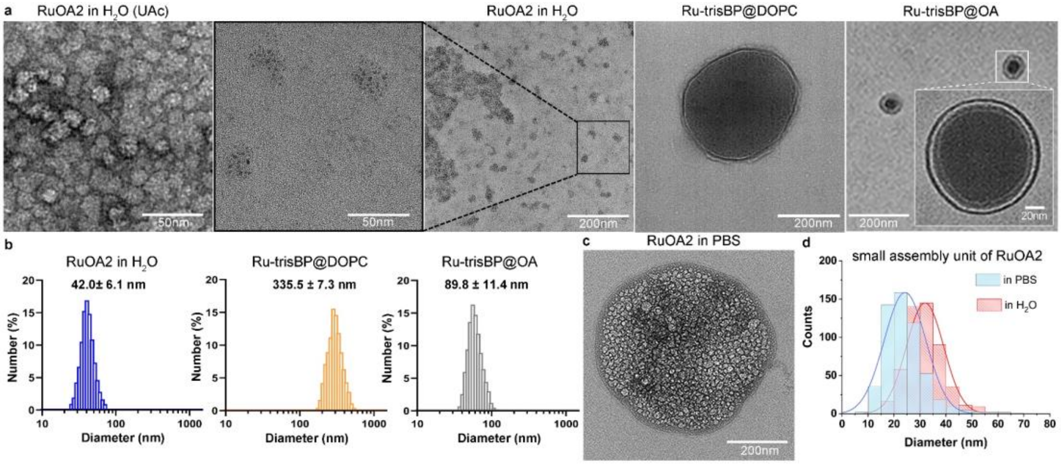


**Figure S3.** (a) TEM images of RuOA2 in water with and without uranyl acetate (UAc) staining, Ru-trisBP@DOPC and Ru-trisBP@OA in water without negative staining. (b) Particle size distribution profiles of RuOA2 ruthenosomes, Ru-trisBP@DOPC, and Ru-trisBP@OA liposomes in water. (c) TEM images of RuOA2 ruthenosomes in PBS buffer, acquired using a Talos-L120C transmission electron microscope. (d) Size distribution profiles of smaller vesicle-like structures within RuOA2 ruthenosomes and vesicles formed by RuOA2 after negative staining, calculated through imaging analysis.

| Cell | IC50 (μM) |
|---|---|
| CT26 | 0.26±0.01 |
| HCT116 | 0.33±0.008 |
| H1975 | 3.73±0.43 |
| MCF7 | 4.11±0.13 |
| Hela | 10.43±0.47 |
| HepG2 | 16.33±0.88 |
| PANC-1 | 16.75±1.48 |
| A549 | 17.08±1.78 |
| HEK 293T | 20.27±2.26 |

**Figure S4.** IC50 values of RuOA2 against various cancer cell lines. Non-linear regression analysis was performed using GraphPad Prism software to calculate the IC50 values from the dose-response curves.

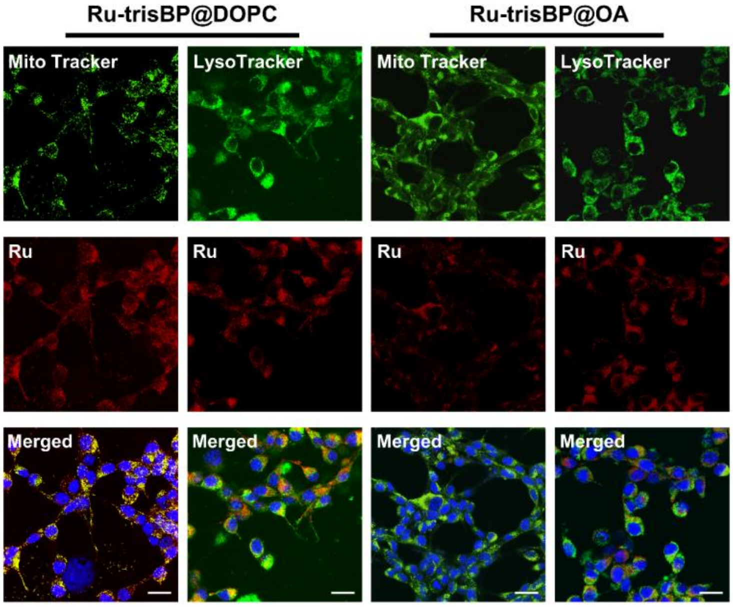


**Figure S5.** CLSM images of HCT116 cells incubated with RutrisBP@DOPC or RutrisBP@OA for 4 hours as well as mitochondria tracker green and lysosome tracker green for 30 minutes. Scale bars represent 20 μm.

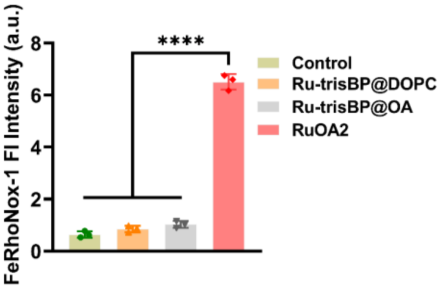


**Figure S6.** Quantification of luminescence intensity of FeRhoNox-1 from the microscopy images shown in Figure 4a. Data are presented as mean ± SD (n=3). ***$p < 0.001$.

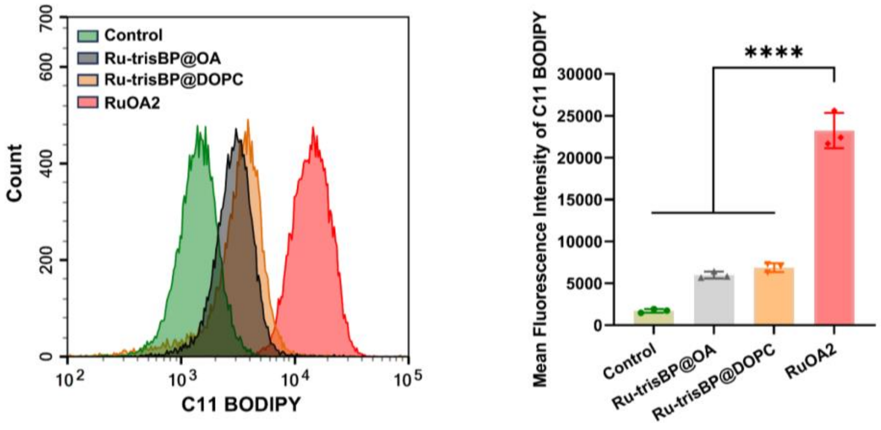


**Figure S7.** Flow cytometry (FCM) analysis of lipid peroxidation (LPO) levels in HCT116 cells treated with various formulations (300 nM) for 24 hours, detected using the C11-BODIPY probe (left panel). Quantification of C11-BODIPY fluorescence intensity (right panel). Data are presented as Mean ± SD (n=3).

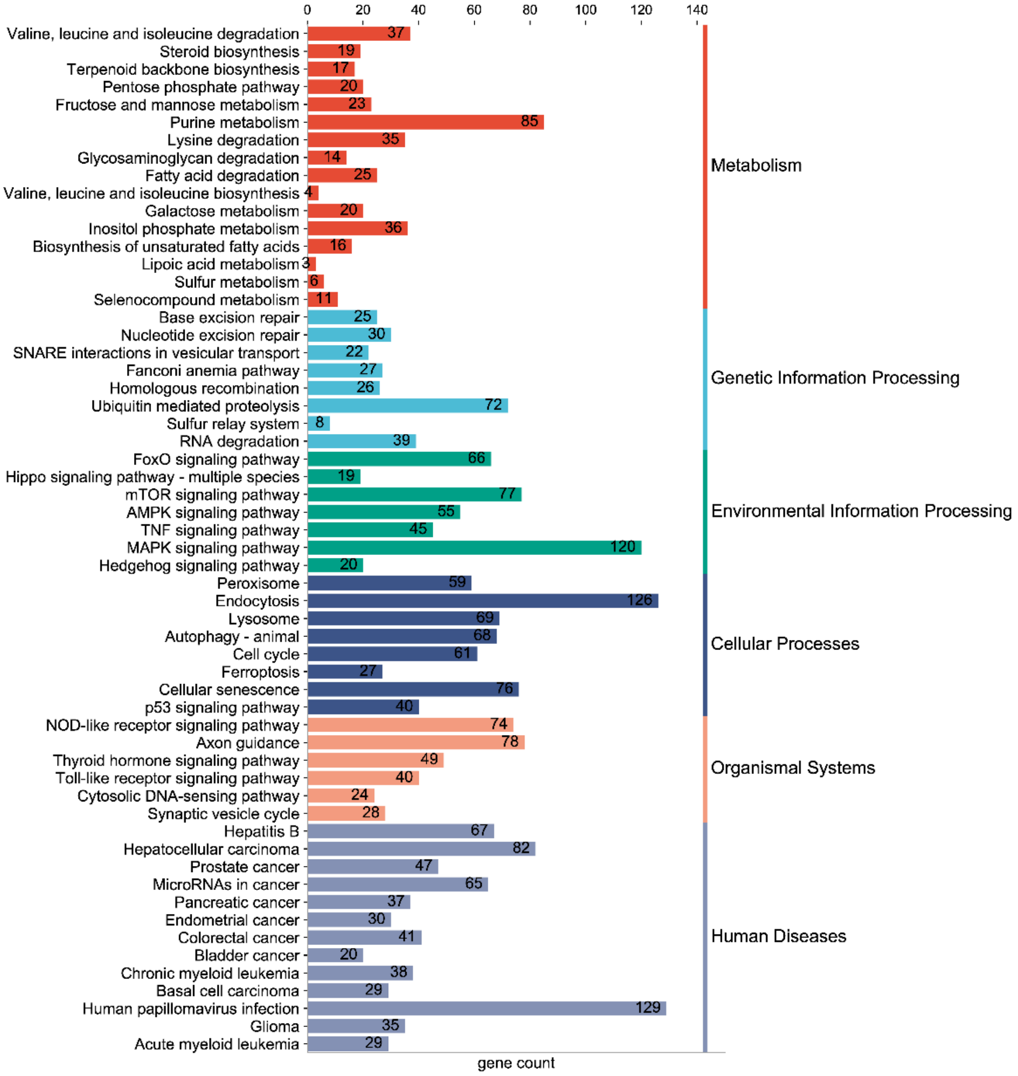


**Figure S8.** All clusters in KEGG pathway enrichment analysis of differentially expressed genes (DEGs) after **RuOA2** (300 nM, 24 hours) treatment.

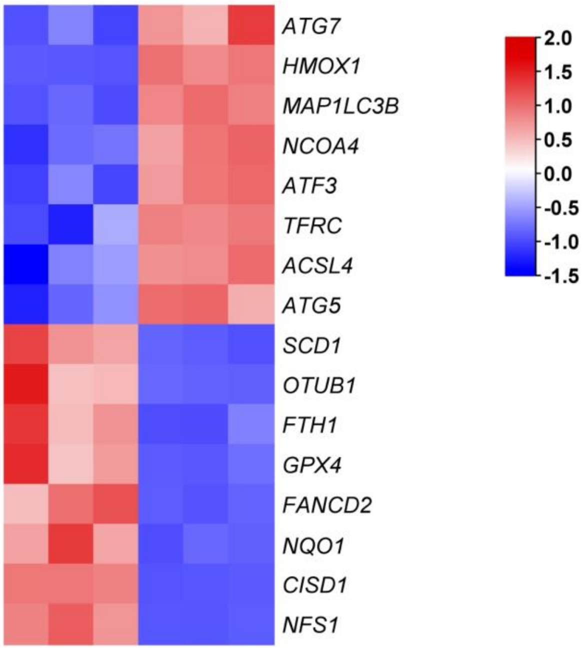


**Figure S9.** Heat map of gene expression before (left) and after (right) **RuOA2** (300 nM, 24 h) treatment.

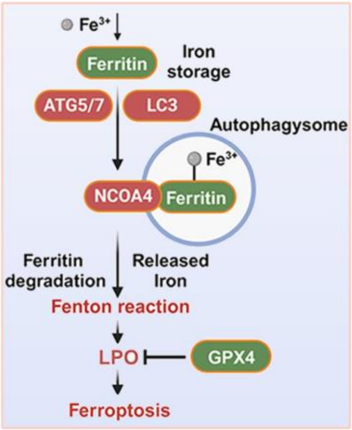


**Figure S10.** Pathways involving differentially expressed genes (DEGs) associated with autophagy and ferritinophagy. Red boxes indicate upregulated genes, while green boxes indicate downregulated genes.

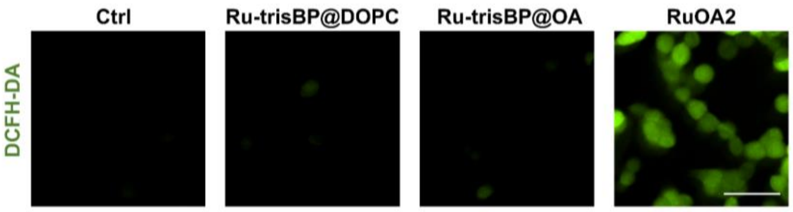


**Figure S11.** Fluorescent images of HCT116 cells incubated with RuOA2 for 4 hours as well as the ROS scavenger DCFH-A for 30 min. Scale bar represents 50 μm.

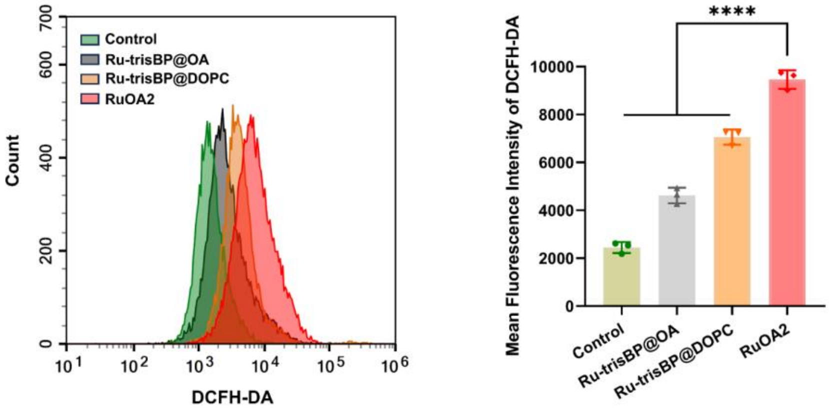


**Figure S12.** FCM analysis of ROS content in HCT116 cells treated with various formulations (300 nM) for 24 hours, using DCFH-A probe detection (left panel). Quantification of DCFH-A fluorescence intensity in left panel (right panel). Data presented as Mean ± SD (n=3).

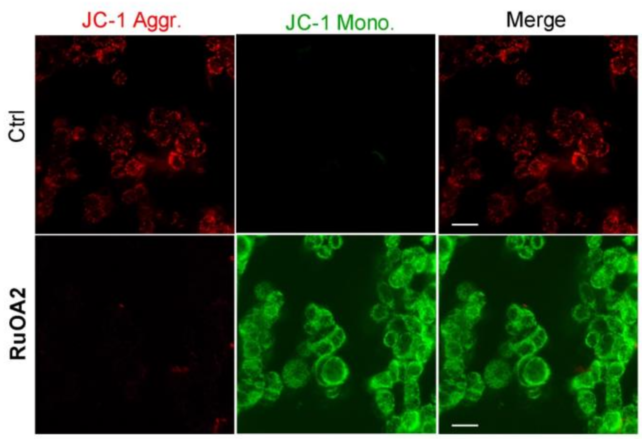


**Figure S13.** HCT116 cells stained with JC-1 showing the loss of red-fluorescent J-aggregate at hyperpolarized membrane potentials and an increased green-fluorescent monomer at depolarized membrane upon the treatment of RuOA2 (1μM) for 4 hours. Scale bars represent 20 μm.

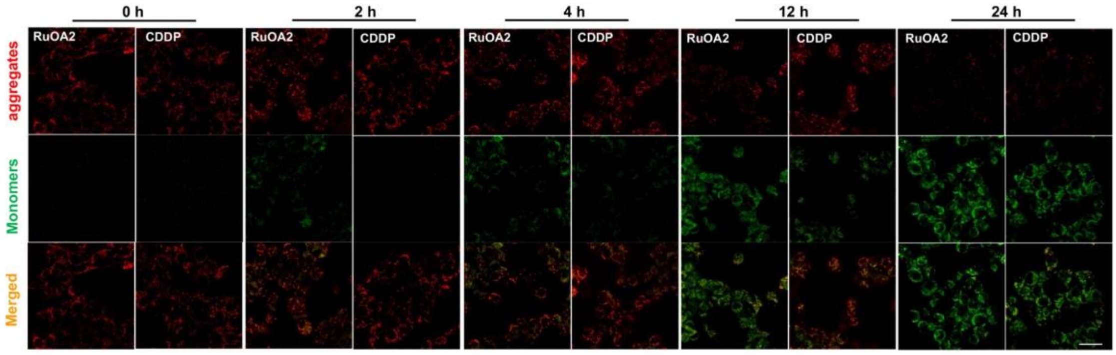


**Figure S14.** Evaluation of mitochondrial membrane potential changes in HCT116 cells treated with 300 nM RuOA2 or 10 μM CDDP, assessed using JC-1 staining at different time points. Scale bar represents 20 μm.

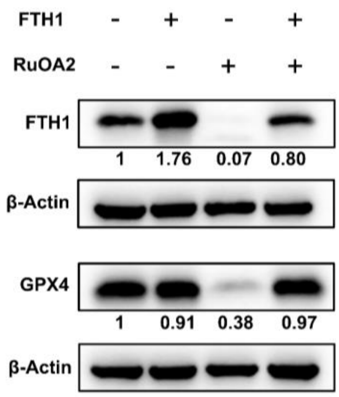


**Figure S15.** Western blotting analysis of FTH1 and GPX4 expression in HCT116 cells transfected with either the control vector or FTH1-overexpressing plasmids, with or without treatment with 300 nM RuOA2 for 24 hours.

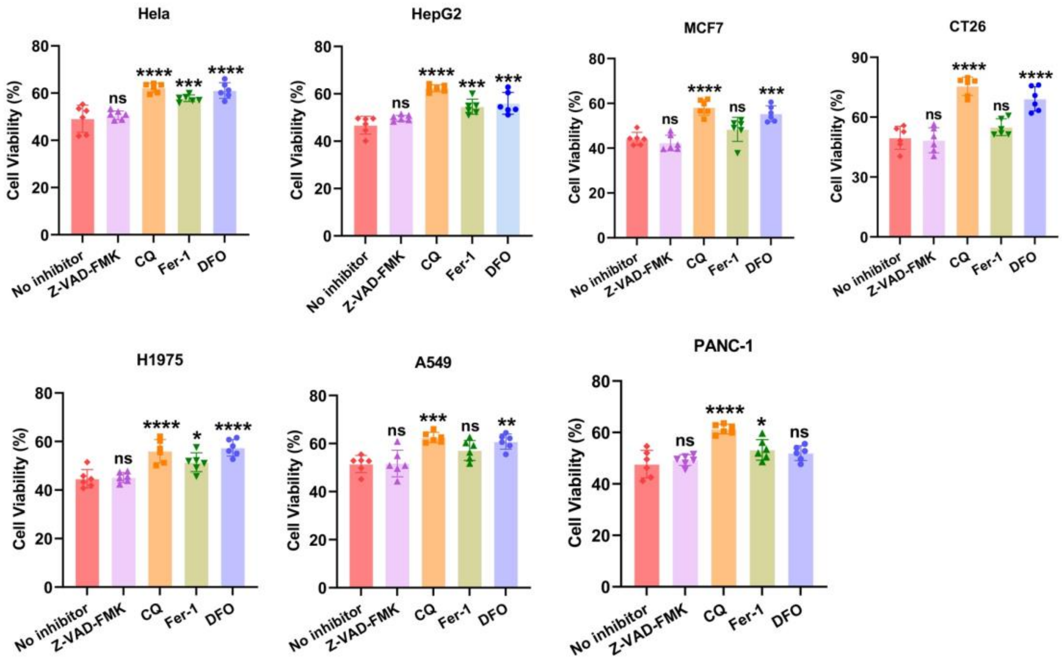


**Figure S16.** A panel of tumor cell lines showing cell viability following 24-hour treatment with RuOA2 at IC50 concentrations, with or without pre-treatment using various PCD inhibitors. Data are presented as Mean ± SD (n=8). ns: not significant; ***p < 0.001, ****p < 0.0001.

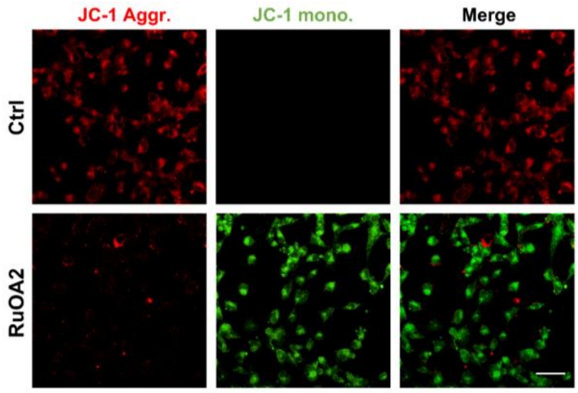


**Figure S17.** CT26 cells stained with JC-1 exhibit a loss of red-fluorescent J-aggregates at hyperpolarized membrane potentials and an increase in green-fluorescent monomers at depolarized membrane potentials following treatment with **RuOA2** (1 μM) for 2 hours. Scale bar represents 50 μm.

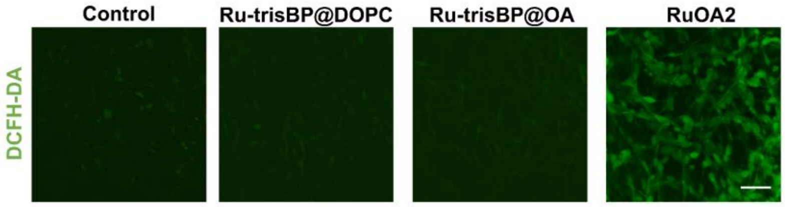


**Figure S18.** Fluorescent images of CT26 cells incubated with RuOA2 (300 nM) for 24 hours, followed by treatment with the ROS scavenger DCFH-A for 30 min. Scale bar represents 50 μm.

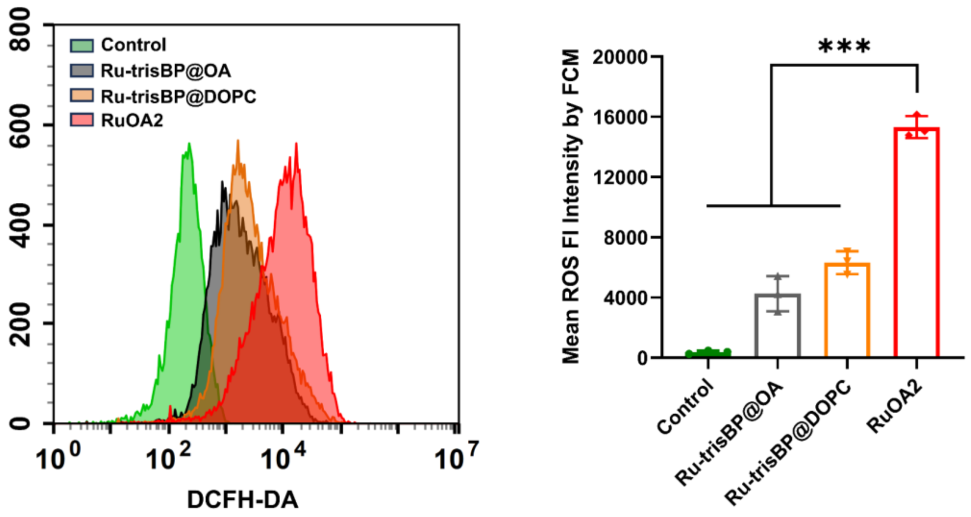


**Figure S19.** FCM analysis of ROS content in CT26 cells treated with various formulations (300 nM) for 24 hours, using DCFH-A probe detection (left panel). Quantification of DCFH-A fluorescence intensity in left panel (right panel). Data presented as Mean ± SD (n=3).

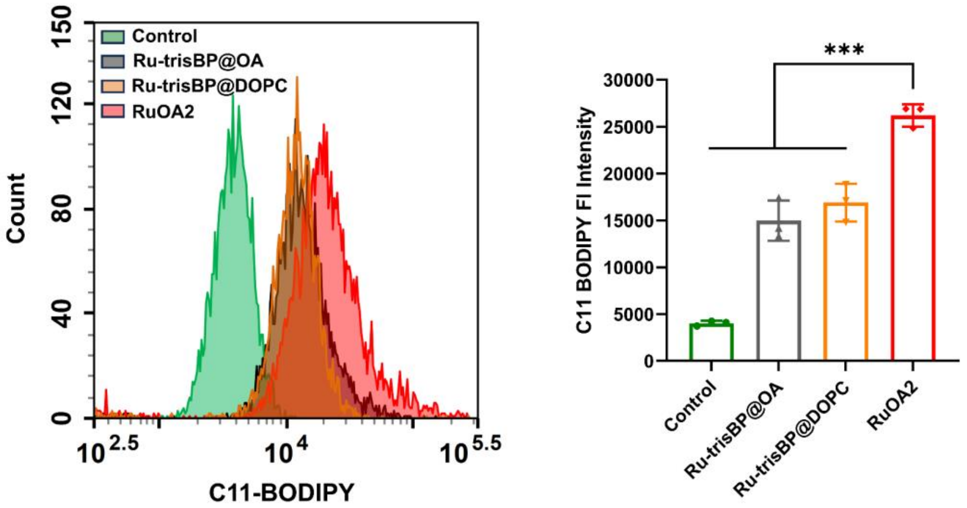


**Figure S20.** Flow cytometry (FCM) analysis of lipid peroxidation (LPO) levels in CT26 cells treated with various formulations (300 nM) for 24 hours, detected using the C11-BODIPY probe (left panel). Quantification of C11-BODIPY fluorescence intensity (right panel). Data presented as Mean ± SD (n=3).

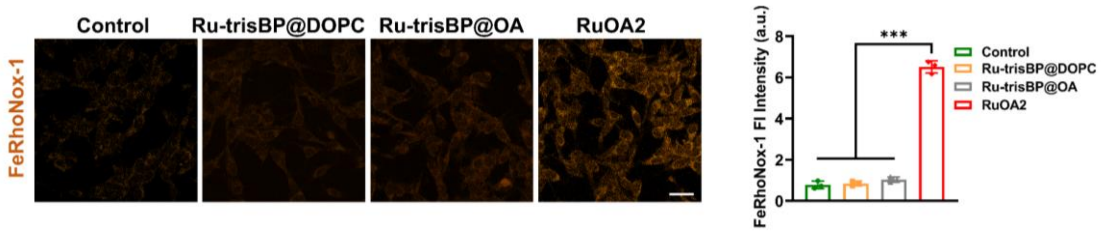


**Figure S21.** Fluorescent images of CT26 cells treated with Ru-complexes (300 nM, 24 hours) and stained with FeRhoNox-1. Scale bar represents 50 μm.

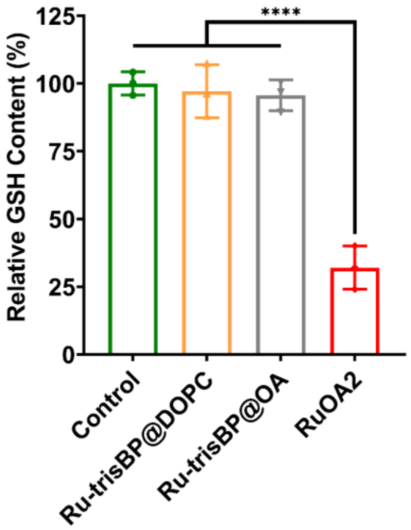


**Figure S22.** Relative intracellular GSH levels in CT26 cells treated with Ru-trisBP@DOPC, Ru-trisBP@OA, and RuOA2 at a concentration of 300 nM for 24 hours. Mean ± SD (n = 3).

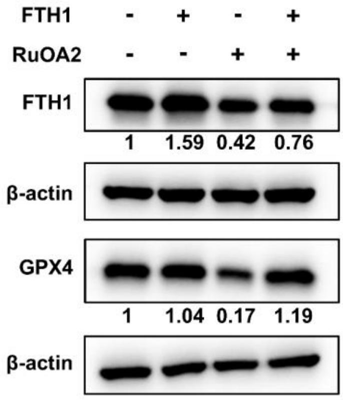


**Figure S23.** Western blot analysis of FTH1 and GPX4 expression in CT26 cells and FTH1-overexpressing CT26 cells, with or without RuOA2 treatment (300 nM) for 24 hours.

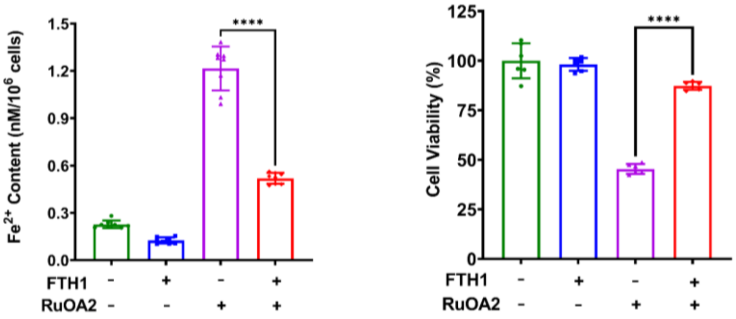


**Figure S24.** Ferrous ion content in CT26 cells transfected with either the control vector or FTH1-overexpressing plasmids, with or without treatment with 300 nM RuOA2 for 24 hours (left panel). Mean ± SD (n=3). Viability of CT26 cells transfected with either the control vector or FTH1-overexpressing plasmids, with or without treatment with 300 nM RuOA2 for 24 hours (right panel). Mean ± SD (n=6).

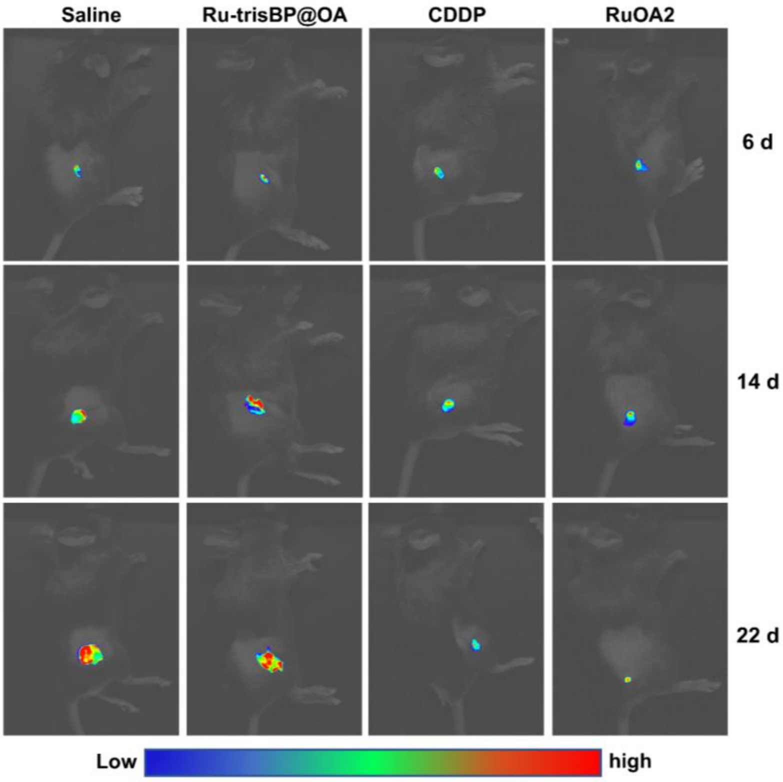


**Figure S25.** Changes in luciferase signals following tumor cell inoculation at 6, 14, and 22 days (n=6).

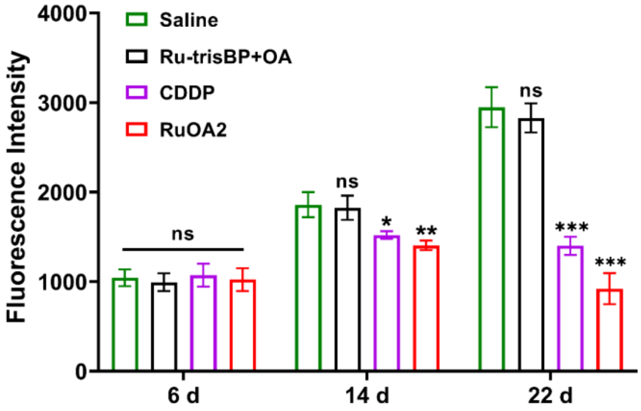


**Figure S26.** Quantification of luciferase fluorescence intensity shown in Figure S25. Mean ± SD (n=6).

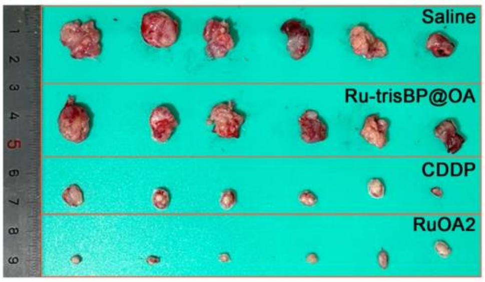


**Figure S27.** Photographs of excised tumors after various treatments.

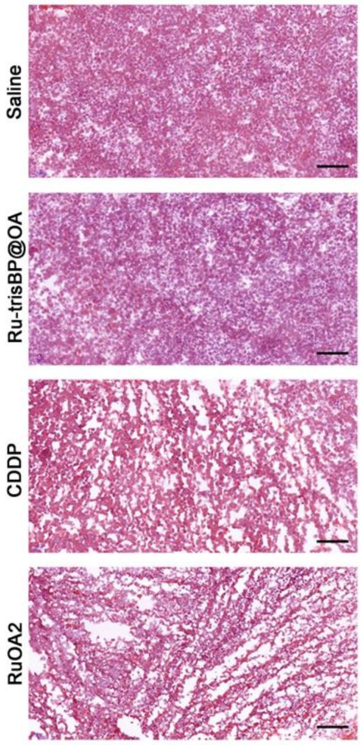


**Figure S28.** H&E staining of tumor tissues after four-dose of various treatments.

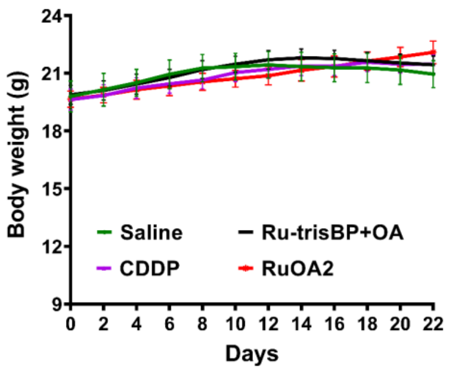


**Figure S29.** Body weight curves of mice under various treatments. Mean ± SD (n=6).

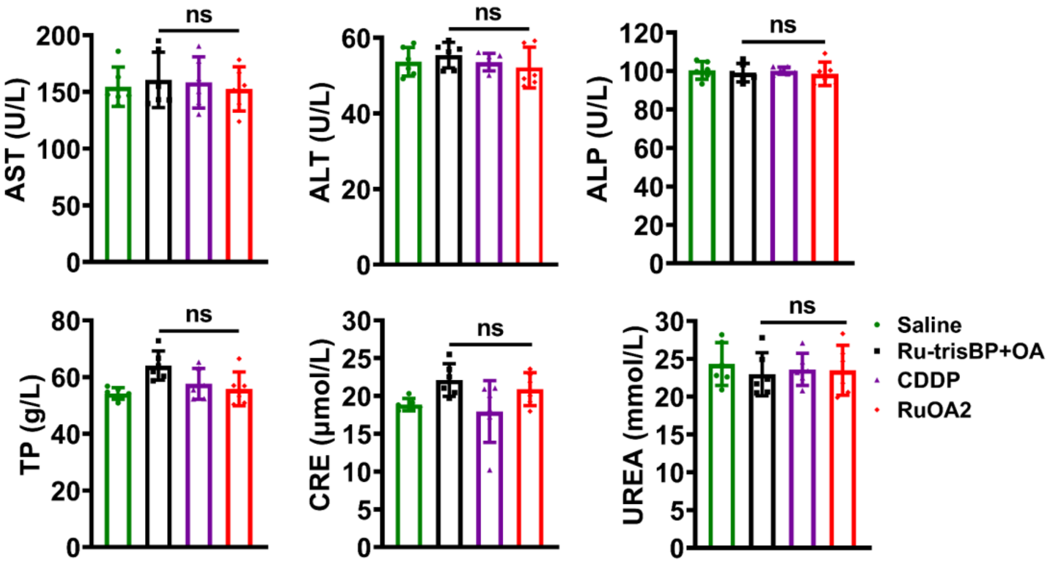


**Figure S30.** Blood biochemical analysis of mice after various treatments. Mean ± SD (n=6). ns: no significant.

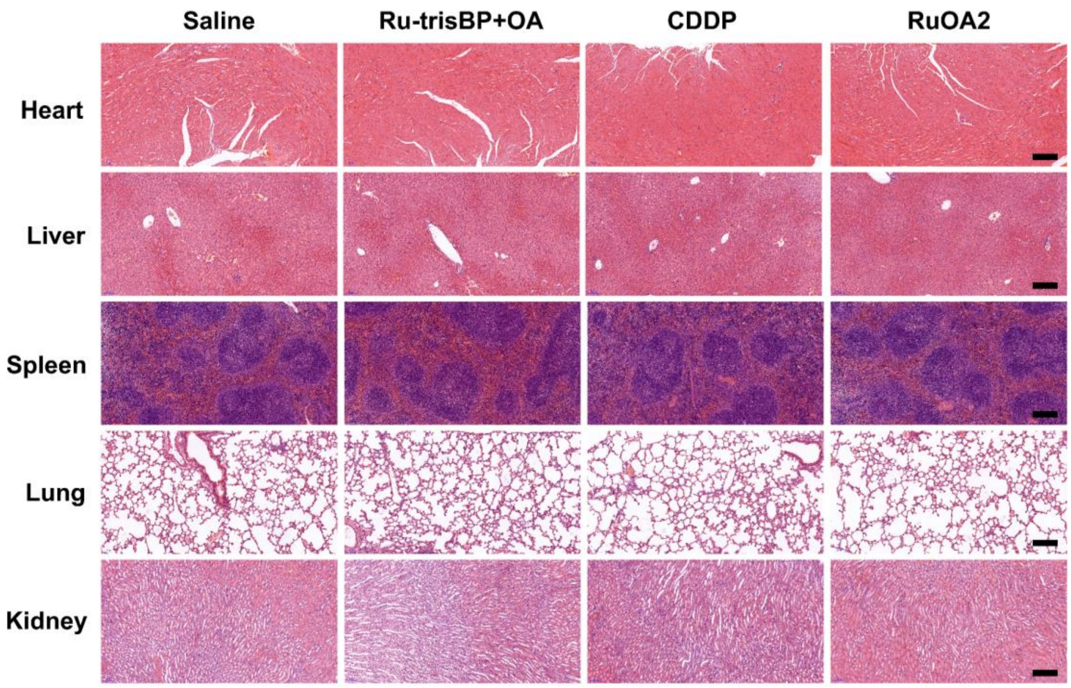


**Figure S31.** H&E staining histopathology results in all groups. Scale bars represent 200 μm.

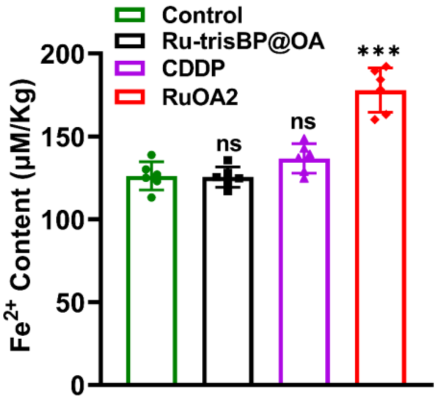


**Figure S32.** $Fe^{2+}$ content in tumor tissues under various treatments. Mean ± SD (n=6). ns: no significant, ***p < 0.001.

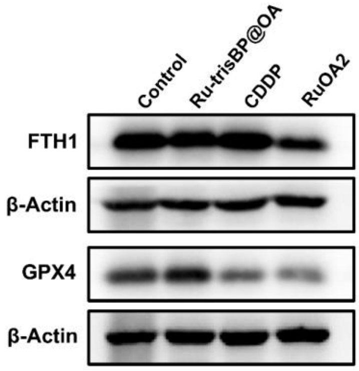


**Figure S33.** Western blotting analysis of FTH1 and GPX4 expression in tumor tissues under various treatments.

**Attachment 1.** $^{1}H$ NMR and $^{13}C$ NMR spectra of **bpy-COOMe** in $CDCl_3$.

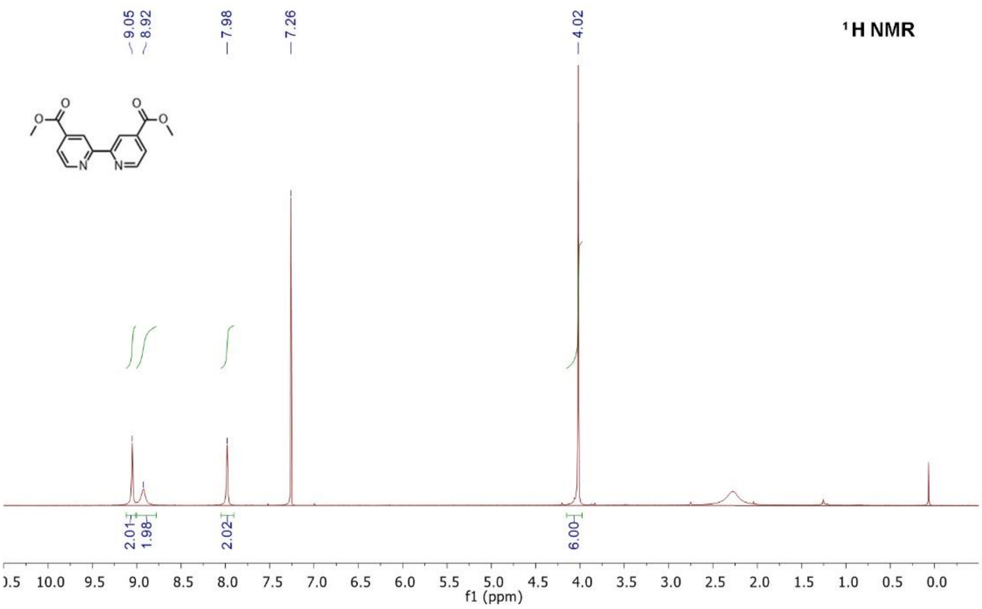


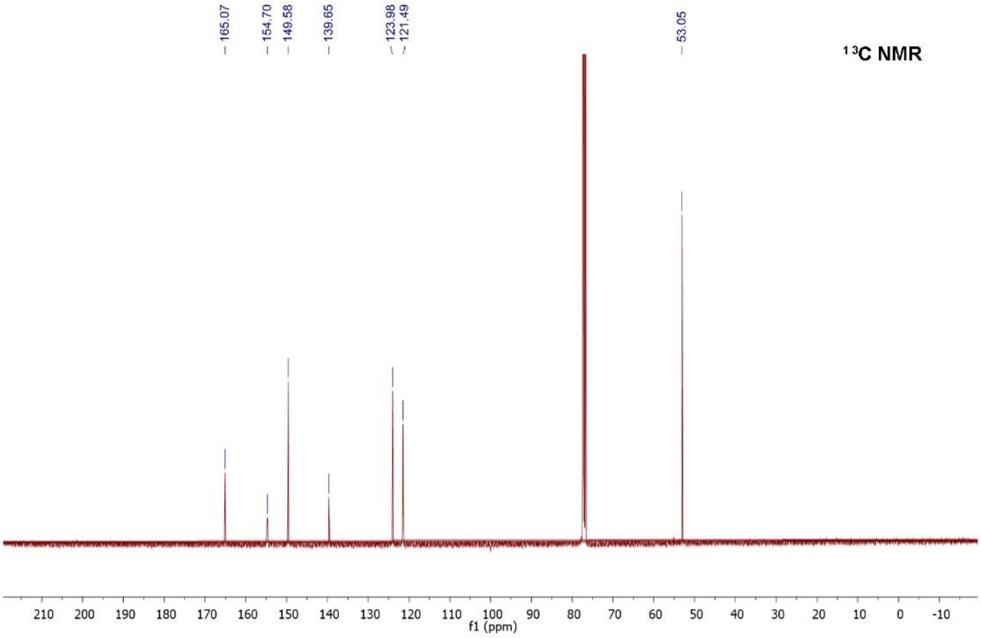

**Attachment 2.** ESI-MS spectra of **Ru-1**.

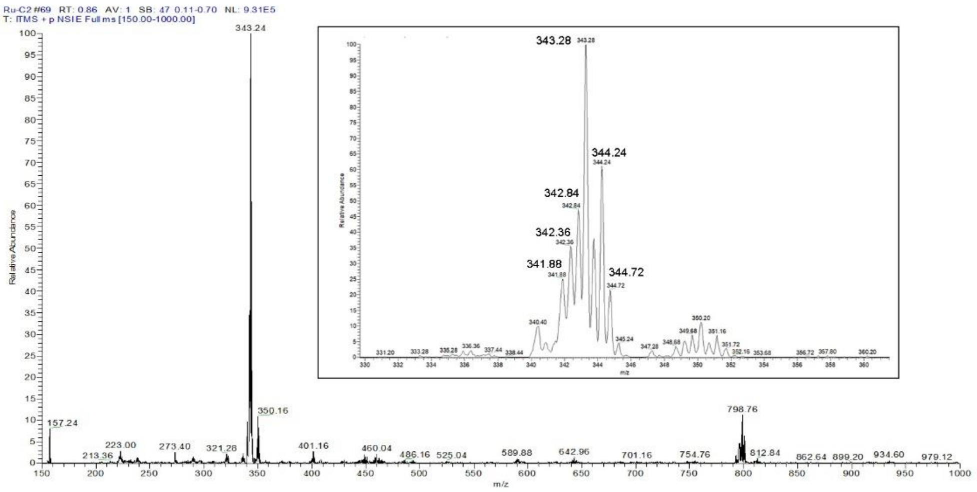

**Attachment 3.** ¹H NMR and ¹³C NMR spectra of **Ru-1** in CDCN.

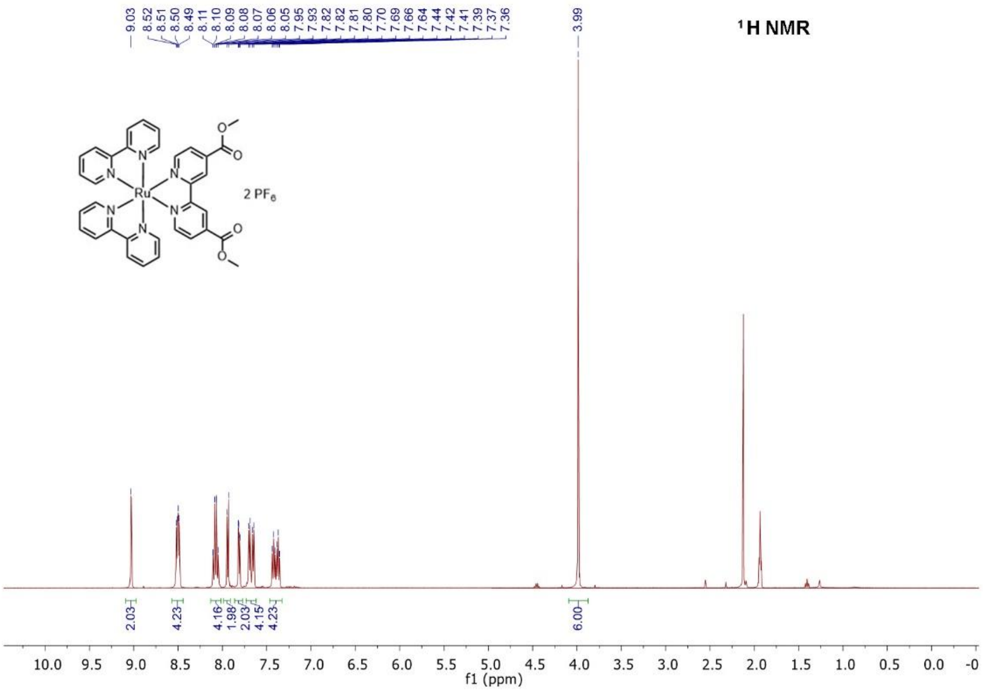


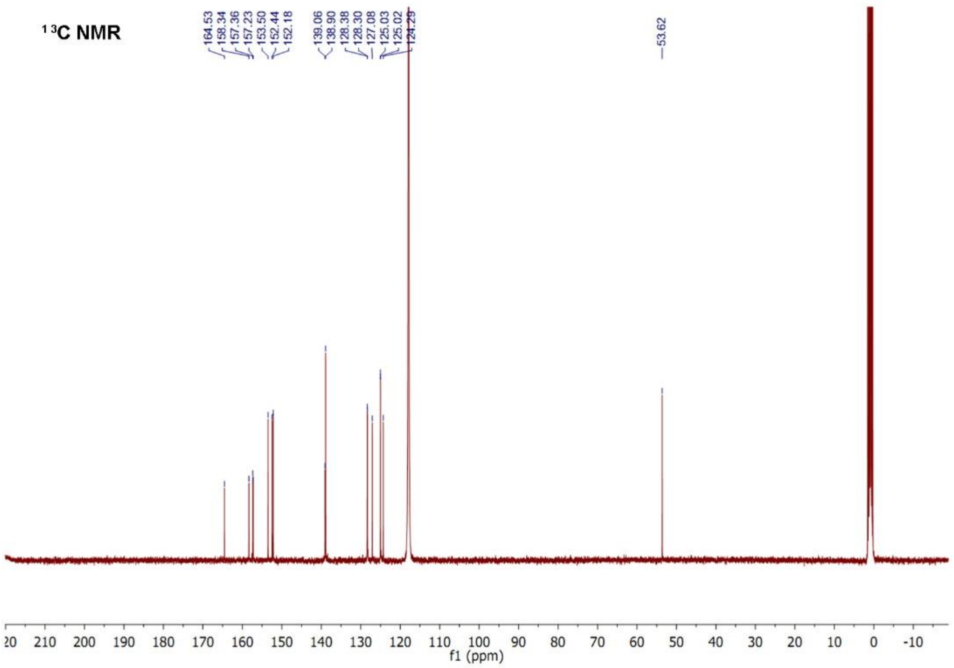

**Attachment 4.** ESI-MS spectra of **Ru-2**.

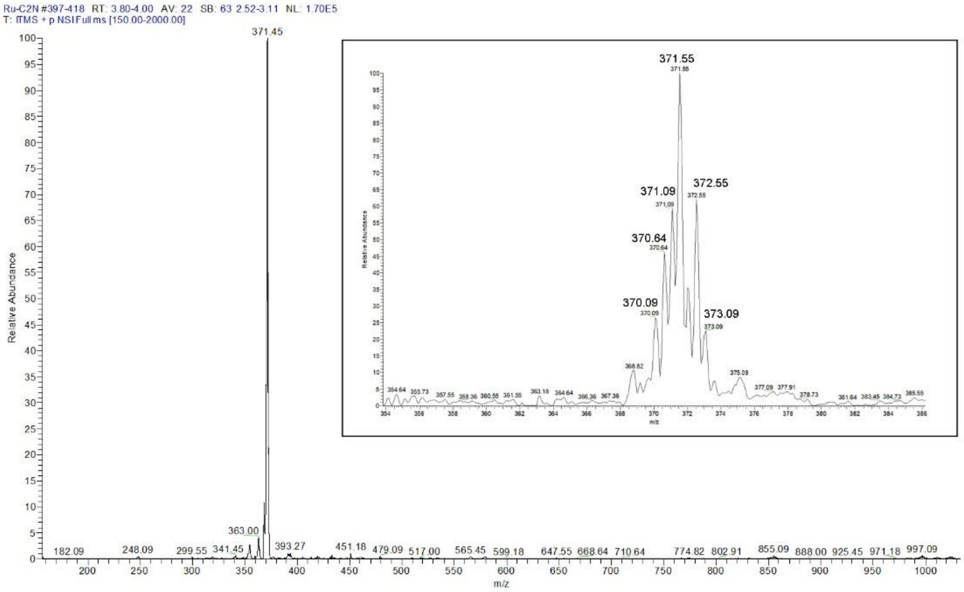

**Attachment 5.**[1]H NMR and [13]C NMR spectra of **Ru-2** in CDCN.

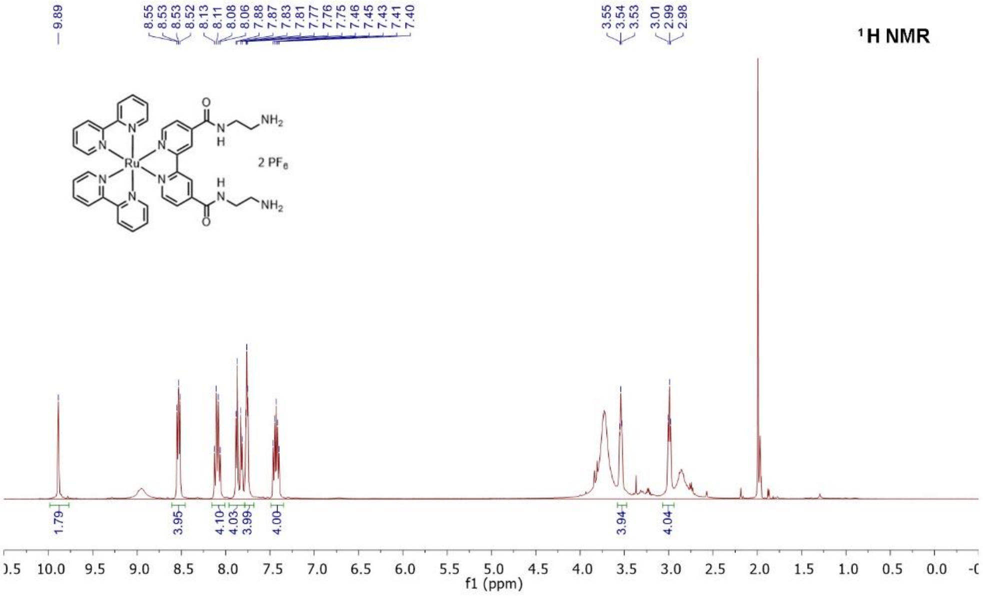


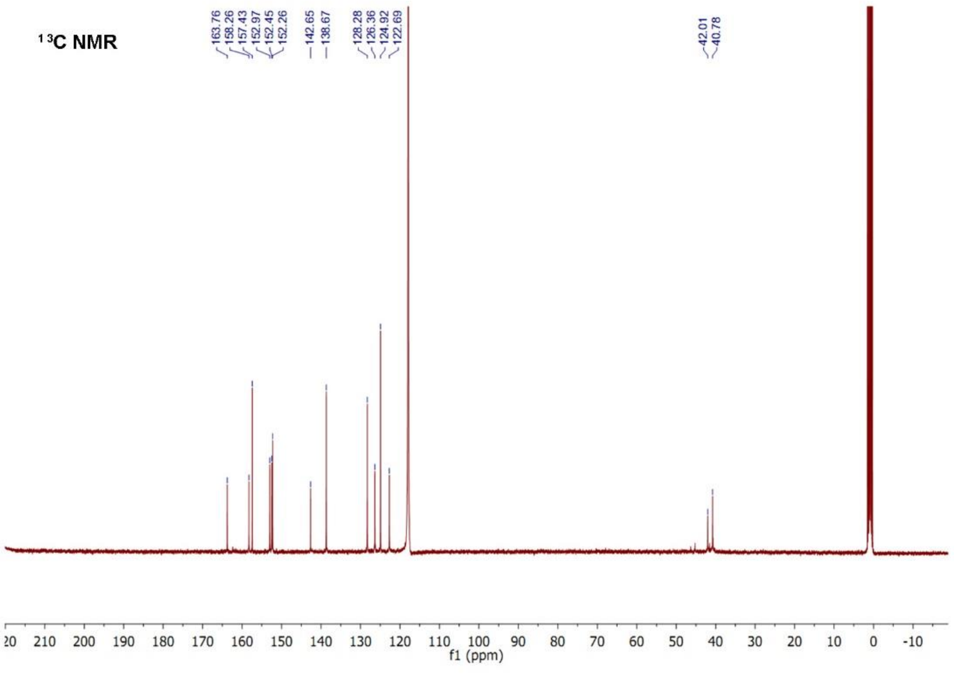

**Attachment 6.** ESI-MS spectra of **RuOA2**.

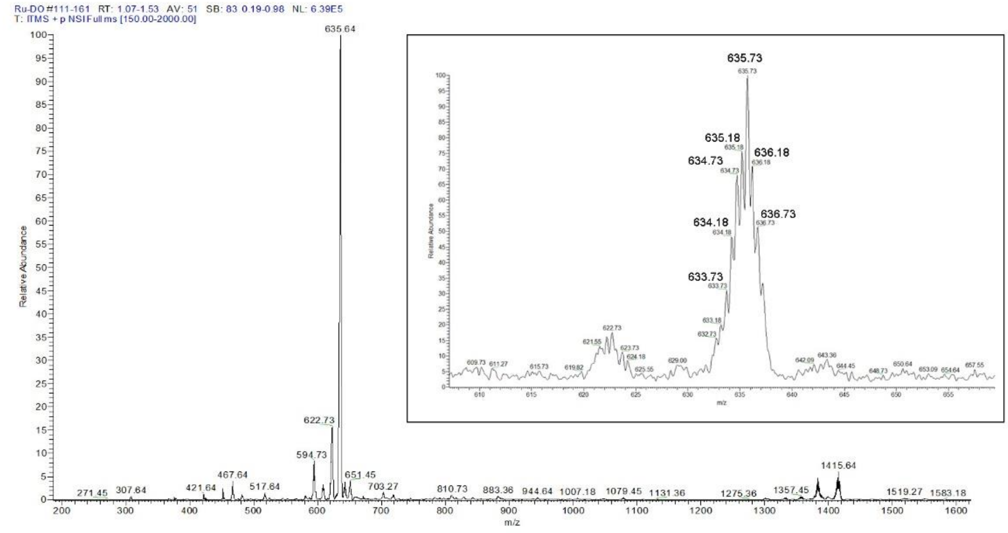

**Attachment 7.** ¹H NMR and ¹³C NMR spectra of **RuOA2** in MeOD.

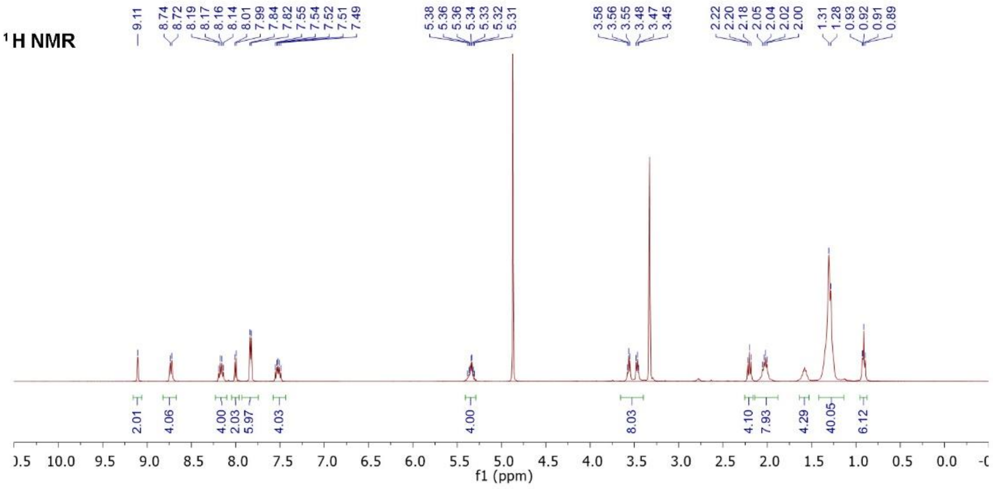


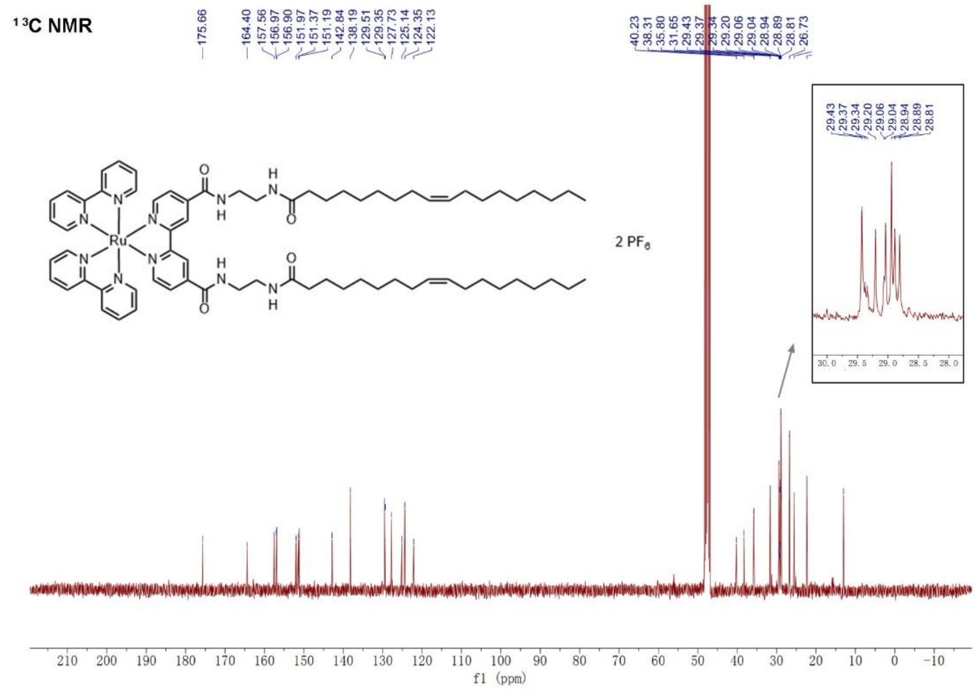

**Attachment 8.** $^{31}P$ NMR and $^{19}F$ NMR spectra of **RuOA2** in MeOD.

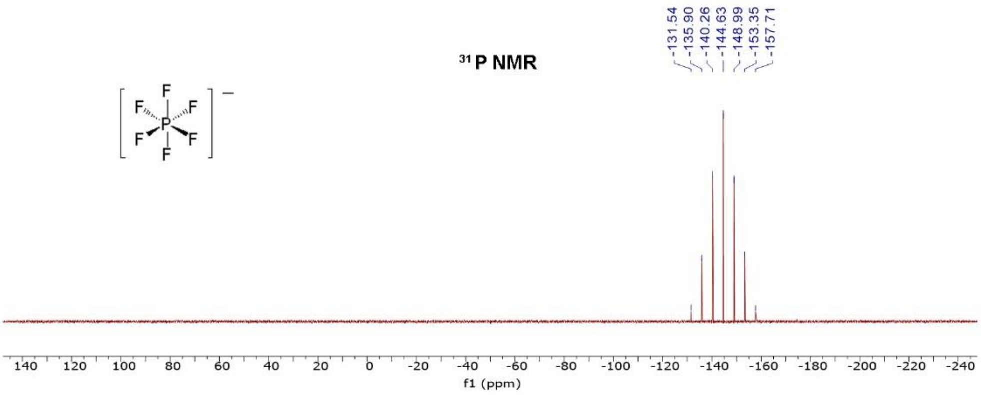


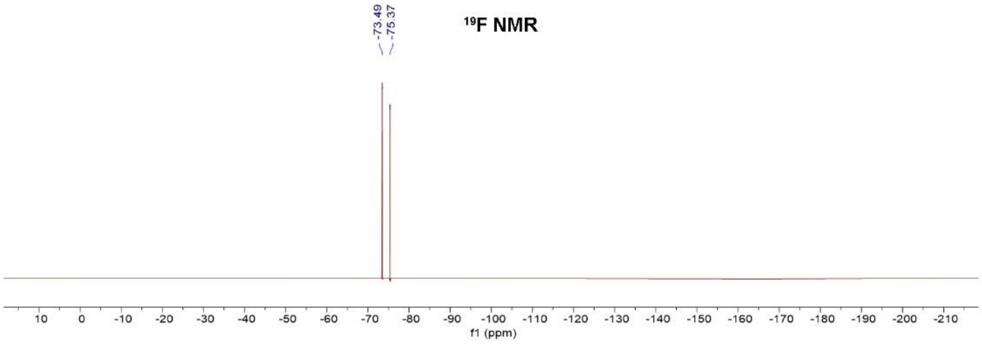
